# SPADIS: An Algorithm for Selecting Predictive and Diverse SNPs in GWAS

**DOI:** 10.1101/256677

**Authors:** Serhan Yilmaz, Oznur Tastan, A. Ercument Cicek

## Abstract

Phenotypic heritability of complex traits and diseases is seldom explained by individual genetic variants identified in genome-wide association studies (GWAS). Many methods have been developed to select a subset of variant loci, which are associated with or predictive of the phenotype. Selecting connected SNPs on SNP-SNP networks have been proven successful in finding biologically interpretable and predictive SNPs. However, we argue that the connectedness constraint favors selecting redundant features that affect similar biological processes and therefore does not necessarily yield better predictive performance. In this paper, we propose a novel method called SPADIS that favors the selection of remotely located SNPs in order to account for their complementary effects in explaining a phenotype. This is achieved by maximizing a submodular set function with a greedy algorithm that ensures a constant factor approximation to the optimal solution. We compare SPADIS to the state-of-the-art method SConES, on a dataset of *Arabidopsis Thaliana* with continuous flowering time phenotypes. SPADIS has better average phenotype prediction performance in 15 out of 17 phenotypes when the same number of SNPs are selected and provides consistent improvements across multiple networks and settings on average. Moreover, it identifies more candidate genes and runs faster. We also investigate the use of Hi-C data to construct SNP-SNP network in the context of SNP selection problem for the first time, which yields improvements in regression performance across all methods. SPADIS is available at http://ciceklab.cs.bilkent.edu.tr/spadis

## I. Introduction

Genome-Wide Association Studies (GWAS) have led to a wide range of discoveries over the last decade where individual variations in DNA sequences, usually single nucleotide polymorphisms (SNPs), have been associated with phenotypic differences (Visscher *et al*., 2017). However, individual variants often fail to explain the heritability of complex traits and diseases (Manolio *et al*., 2009; Goldstein *et al*., 2009) as a large number of variants contribute to these phenotypes and each variant has a small overall effect (Kraft and Hunter, 2009; Christensen and Murray, 2007). Thus, evaluating and associating multiple loci with a given phenotype is critical (Moore *et al*., 2010; Cordell, 2009). Indeed, detecting genetic interactions (epistasis) among pairs of loci has proven to be a powerful approach as discussed in several reviews (Phillips, 2008; Cordell, 2009; Wang *et al*., 2010a; Wei *et al*., 2014).

Detecting higher-order combinations of genetic variations is computationally challenging. For this reason, exhaustive search approaches have been limited to small SNP counts (up to few hundreds) (Nelson *et al*., 2001; Ritchie *et al*., 2001; Lou *et al*., 2007; Lehár *et al*., 2008; Hua *et al*., 2010; Fang *et al*., 2012) and greedy search algorithms have been limited to searching for small combinations of SNPs – mostly around 3 (Storey *et al*., 2005; Evans *et al*., 2006; Yosef *et al*., 2007; Varadan and Anastassiou, 2006; Varadan *et al*., 2006; Zhang and Liu, 2007; Herold *et al*., 2009; Tang *et al*., 2009; Jiang *et al*., 2009; Zhang *et al*., 2010; Wang *et al*., 2010b; Wan *et al*., 2010; Guo *et al*., 2014; Ding *et al*., 2015; Ayati and Koyutürk, 2016; Tuo *et al*., 2017). Multivariate regression-based approaches have been used (Shi *et al*., 2008; Wu *et al*., 2009; Cho *et al*., 2010; Wang *et al*., 2011a; Rakitsch *et al*., 2012). However, (i) their predictive power is limited, (ii) incorporation of biological information in the models is not straightforward, and finally (iii) selected SNP set is often not biologically interpretable (Azencott *et al*., 2013).

Assessing the significance of loci by grouping them based on functionally related genes, such as pathways, reduces the search space for testing associations and leads to the discovery of more interpretable sets (Wang *et al*., 2011b; de Leeuw *et al*., 2015). Unfortunately, using gene sets and exonic regions for association restricts the search space to coding and nearby-coding regions. However, most of the genetic variation fall into non-coding genome (Hindorff *et al*., 2009) and our knowledge of pathways are incomplete.

An alternative strategy to avoid literature bias is to select features on the SNP-SNP networks by applying regression based methods with sparsity and connectivity constraints (Jacob *et al*., 2009; Huang *et al*., 2011). These regularized methods jointly consider all predictors in the model as opposed to univariate test of associations. Nevertheless, using a SNP-SNP interaction network with these regression based methods on GWAS yields intractable number of interactions. An efficient method called SConES uses a minimum graph cut-based approach to select predictive SNPs over a network of hundreds of thousands of SNPs (Azencott *et al*., 2013; Sugiyama *et al*., 2014). In their network, edges denote either (i) spatial proximity on the genomic sequence or (ii) functional proximity as encoded with PPI closeness of loci. The method selects a connected set of SNPs that are individually related to the phenotype under additive effect model and has been shown to perform better than graph-regularized regression-based methods.

We argue that enforcing the selected features to be in close proximity encourages the algorithm to pick features that are in linkage disequilibrium or that have similar functional consequences. One extreme choice of this approach would be to choose all SNPs that fall into the same gene if they are individually found to be significantly associated with the phenotype. When there is an upper limit on the number of SNPs to be selected, this leads to selecting functionally redundant SNPs and miss variants that cover different processes. Genetic complementation, on the other hand, is a well-known phenomenon where multiple loci in multiple genes need to be mutated in order to observe the phenotype (Fincham, 1968). While there are numerous examples of long-range (trans) genetic interactions for transcription control (Miele and Dekker, 2008) and long-range epistasis is evident in complex genetic diseases such as type 2 diabetes (Wiltshire *et al*., 2006), such complementary effects may not be treated with this approach. For disorders with complex phenotypes like Autism Spectrum Disorder (ASD), this would be even more problematic since multiple functionalities (thus gene modules in the network) are required to be disrupted for an ASD diagnosis, whereas damage in only one leads to a more restricted phenotype (Geschwind, 2008).

We hypothesize that diversifying the SNPs in terms of location would result in *covering* complementary modules in the underlying network that cause the phenotype. Based on this rationale, here, we present SPADIS, a novel SNP selection algorithm over a SNP-SNP interaction network that favors (i) loci with high univariate associations to the phenotype and (ii) that are diverse in the sense that they are far apart on a loci interaction network. In order to incorporate these principles, we design a submodular set scoring function and select SNPs by maximizing this set function. To maximize this set function, we use a greedy algorithm that is guaranteed to return a solution which is a constant factor (1 − 1/*e*) approximate to the optimal solution. We compare our algorithm to the state-of-the-art method SConES, on a GWAS of *Arabidopsis Thaliana (AT)* with 17 continuous phenotypes related to flowering time (Atwell *et al*., 2010). We show that SPADIS has better average regression performance in 15 out of 17 phenotypes with better runtime performance. Moreover, our method always identifies more candidate genes (up to 50%) and always hits more Gene Ontology (GO) terms (up to 20%) on average, indicating that selection of SPADIS is more diverse.

Finally, we employ Hi-C data in the context of SNP selection problem for the first time. Emerging evidence suggests that the spatial organization of the genome plays an important role in gene regulation (Bickmore, 2013) and contacts in 3D have been shown to affect the phenotype (Martin *et al*., 2015; Jäger *et al*., 2015). Hi-C technology detect the can 3D conformation genome-wide and yield contact maps which show loci that reside nearby in 3D (van Berkum *et al*., 2010). We construct a SNP-SNP network based on genomic contacts in 3D as captured by Hi-C and use this network to guide SNP selection. Our results show that use of Hi-C based network provides a slight overall increase in the prediction performance for all methods tested.

## II. Methods

The problem is formalized as a feature selection problem over a network of SNPs. Let *n* be the number of SNPs. The problem is to find a SNP subset *S* with cardinality at most *k* ≪ *n* that explains the phenotype, given a background biological network *G*(*V, E*). In *G*, vertices represent SNPs and edges link loci which are related based on spatial or functional proximity as explained in sections below. *G* can be a directed or an undirected graph.

We utilize a two-step approach. In the first step, we assess the relation of each SNP to the phenotype individually using the Sequence Kernel Association Test (SKAT) (Wu *et al*., 2011). In the second step, our goal is to maximize the total score of SNP set while ensuring the selected set consists of SNPs that are remotely located on the network. Under the additive effect model, we define the set function shown in Equation 1 to encode this intuition.

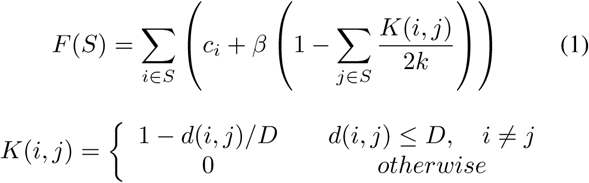

Here **c** is the scoring vector such that 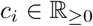 indicates the level of the *i*-th SNP’s association with the phenotype. 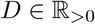 is a distance limit parameter and *d*(*i, j*) is the shortest path between vertices *i, j* ∈ *V*. Note that, *d*(*i, j*) = ∞ if *j* is not reachable from *i*. *K*(*i, j*) is a function that penalizes vertices that are in *close* proximity. That is, the vertices *i* and *j* are considered *close* if and only if *d*(*i, j*) ≤ *D*. The second parameter, 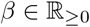 controls the penalty to be applied when two close vertices are jointly included in *S*. Note that, *K*(*i, j*) ∈ [0, 1], ∀*i*, *j* ∈ *V* and *c_i_* is non-negative.

Our aim is to find a subset of SNPs *S*\* of size *k* that maximizes *F*:

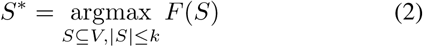

Subset selection problem with cardinality constraint is NP-hard. Thus, exhaustive search is infeasible when *k* or *V* is not small. We make use of the fact that the function defined in Equation 1 is submodular. Although submodular optimization itself is NP-hard as well (Krause and Guestrin, 2005), the greedy algorithm given in **Algorithm 1**, proposed by Nemhauser *et al*. (1978), guarantees a 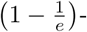 factor approximation to the optimal solution under cardinality constraint for monotonically non-decreasing and non-negative submodular functions. The greedy algorithm starts with an empty set and at each step, adds an element that maximizes the set function. Note that, this is equivalent to adding elements with the largest marginal gain.

### Algorithm 1 Greedy Algorithm

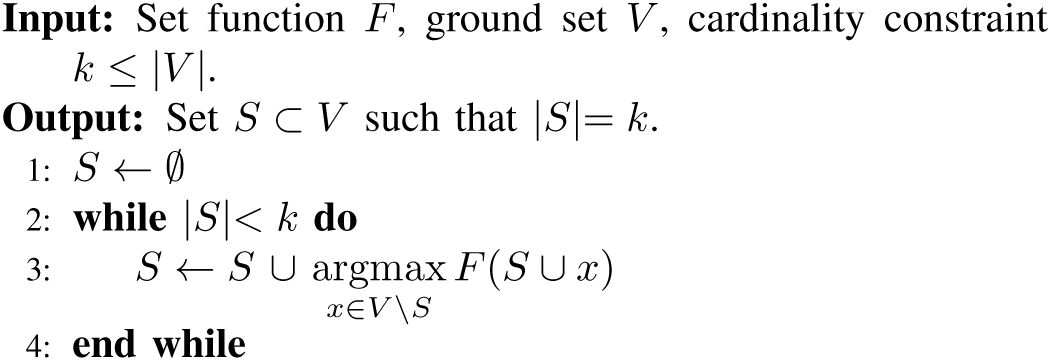

For each of the *k* iterations in the algorithm, where *k* is the size of *S*\*, a single source shortest path problem needs to be solved. Hence, the worst-case time complexity of the algorithm is *O*(*k*(*V* + *E*)) assuming that all edge weights are positive. For undirected graphs, *K*(*i, j*)= *K*(*j, i*) and computations can be reduced by half.

A submodular function is a set function for which the gain in the value of the function after adding a single item decreases as the set size grows (diminishing returns). Next, we prove that *F* is a submodular set function.

### Definition 1.

*V* is the ground set, 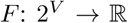 and *S* ⊆ *V*. The marginal gain of adding one element to the set *S* is: *G*(*S, x*) = *F*(*S* ∪ {*x*}) − *F*(*S*) where *x* ∈ *V* \ *S*.

By plugging the definition of *F* in Equation 1, we can rewrite *G*.

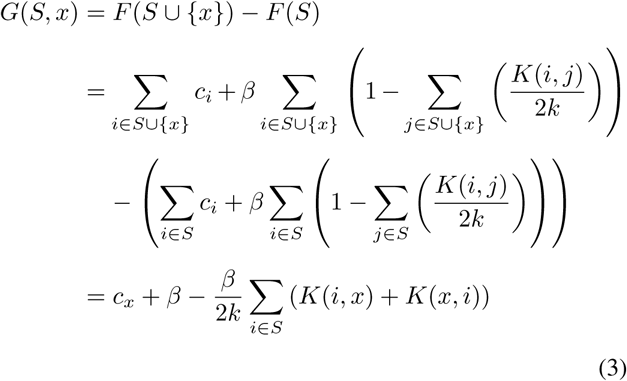

### Definition 2.

A function *F* that is defined on sets, is *submodular* if and only if *G*(*A, x*) ≥ *G*(*B, x*) or equivalently *F*(*A* ∪ {*x*}) − *F*(*A*) ≥ *F*(*B* ∪ {*x*}) − *F*(*B*) for all sets *A, B* where *A* ⊂ *B* ⊂ *V* and *x* ∈ *V* \ *B*.

#### Lemma 1.

*F*(*S*) given in Equation 1 is *submodular*.

##### Proof.

*F* is *submodular* if and only if the following is true:

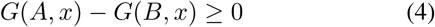

Let *H*(*A, B, x*) be,

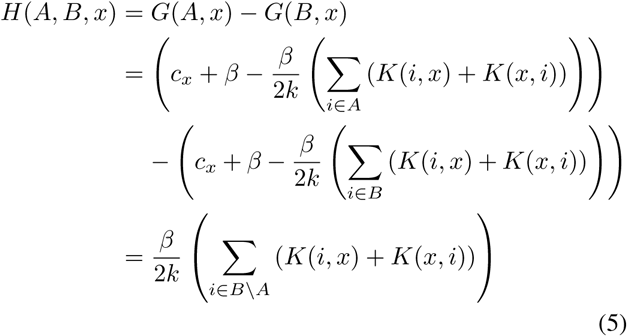

Since *K*(*i, j*) ≥ 0 ∀*i, j* ∈ *V*, *H*(*A, B, x*) ≥ 0. Hence, *F* is *submodular*.

To be able to use the greedy algorithm, *F* must be a monotonically non-decreasing and non-negative function. Below, we prove that *F* satisfies these properties.

### Definition 3.

*F*(*S*) is *monotonically non*-*decreasing* function for sets if and only if the corresponding gain function is always non-negative i.e. *G*(*S, x*) ≥ 0 for all sets *S* ⊂ *V* and *x* ∈ *V*.

#### Lemma 2.

*F*(*S*) given in Equation 1 is *monotonically non*-*decreasing* for sets for which |*S*| ≤ *k*.

##### Proof.

Since *K*(*i, j*) ≤ 1 ∀*ij*, *G*(*S, x*) is bounded such that;

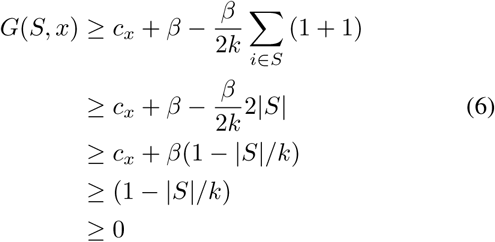

Since |*S*| ≤ *k*, *F*(*S*) is *monotonically non*-*decreasing*.

#### Lemma 3.

*F*(*S*) given in Equation 1 is non-negative for sets |*S*| ≤ *k*.

##### Proof.

For any set *S* = {*v*_1_, *v*_2_, …, *v_n_*} with cardinality *n*, let *S^i^* denote the subset of *S* that contains elements up to the *i*-th element, i.e. *S^i^* = {*v*_1_, *v*_2_, …, *v_i_*} and *S^i^* = ∅ for *i* = 0. *F*(*S*) can be decomposed as the summation of marginal gain functions:

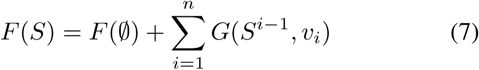

*F*(∅)=0 by the definition of *F*(*S*). **Lemma 2** states that *G*(*S, x*) ≥ 0 for all sets *S* ⊂ *V* and *x* ∈ *V* \ *S* when |*S*| ≤ *k*. Hence, *F*(*S*) ≥ 0 for all sets *S* ⊂ *V* where |*S*| ≤ *k*.

## III. Results

### A. Dataset

We use *AT* genotype and phenotype data from Atwell *et al*. (2010). The dataset includes 17 phenotypes related to flowering times (up to *m* = 180 samples and *n* = 214 051 SNPs). Gene-gene interaction network is constructed based on TAIR protein-protein interaction (PPI) data^1^. SNPs with a minor allele frequency (MAF) < 10% are disregarded (*n* = 173 219 SNPs remained) and population stratification is corrected using the principal components of the genotype data (Price *et al*., 2006). Candidate genes pertaining to each phenotype is retrieved from Segura *et al*. (2012) and used for validating the models. Gene Ontology (GO) annotations are obtained from TAIR (Berardini *et al*., 2004). We obtain the Hi-C data for *AT* from Wang *et al*. (2015) and process the intra-chromosomal contact matrices using the Fit-Hi-C method (Ay *et al*., 2014).

### B. Networks

We construct four undirected SNP-SNP networks. To be able to compare the performances of SPADIS and SConES in a controlled setting, we use three networks defined in Azencott *et al*. (2013): The *GS (gene sequence) network* links loci that are adjacent on the DNA sequence. The *GM (gene membership) network* additionally links two loci if both loci fall into the same gene or they are both close to the same gene below a threshold of 20 000 bp. The *GI (gene interaction) network* also links any two loci if their nearby genes are interacting in the protein interaction network. Note that, GS ⊂ GM ⊂ GI. To investigate the usefulness of the 3D conformation of the genome in this setting, we introduce a new network, GS-HICN which connects loci that are close in 3D in addition to 2D (GS). That is, an edge is added on top of the GS network for loci pairs that are significantly close in 3D (FDR adjusted p-value ≤ 0.05). All networks contain 173 219 vertices. The number of (undirected) edges are as follows: GS: 173 214, GM: 11 661 166, GI: 18 134 516, GS-HICN: 2 919 607.

### C. Compared Methods

We compare SPADIS with the following methods:

*SConES:* A network-constrained SNP selection method with a max-flow based solution (Azencott *et al*., 2013).

*Univariate:* We run univariate linear regression and select SNPs that are found to be significantly associated with the phenotype (FDR-adjusted p-value ≤ 0.05) (Yekutieli and Benjamini, 1999). If the number of SNPs found to be associated is larger than a cardinality constraint of *k* (the maximum number of SNPs to be selected), only the most significant *k* SNPs are picked.

*Lasso:* The Lasso regression (Tibshirani, 1996) that minimizes the prediction error with the *ℓ*1-regularizer of the coefficient vectors. We use the SLEP implementation (Liu *et al*., 2009). *GraphLasso and GroupLasso:* We also compare our method to GraphLasso and GroupLasso (Jacob *et al*., 2009) through simulations, using the implementation in the SLEP package. Due to the prohibitive runtimes of these algorithms, they are excluded from the comparison on *AT* dataset (see *Time Performance* section). For GraphLasso, SNP pairs connected with an edge constitute a separate group, i.e. one such group is constructed for every edge in the network. For GroupLasso, the groups are defined as follows. For *GS*: every consecutive SNP pair on the genome constitute a single group. This is equivalent to setting a group for an edge. For GM: the SNPs *near* (< 20 kbp) a gene are considered as a group, and a separate group is constructed for every gene. For GI: the SNPs that are near interacting genes in the PPI network are combined and formed a single group. The SNPs that are near genes that do not participate in the gene interaction network are assigned to groups based on their gene membership as in GM. For GS-HICN: SNP pairs connected with an edge is considered as a separate group similar to the groups in GraphLasso.

### D. Experimental Setup

A fair comparison among such a diverse range of methods is challenging. SPADIS operates with a cardinality constraint, whereas other methods have parameters that affect the number of selected SNPs. To account for such differences, we compare the methods using either of the following constraints: (1) Tight cardinality constraint where all methods select a fixed number of SNPs which is *k*, and (2) maximum cardinality constraint where the methods are allowed to select SNP sets of different sizes as long as the set sizes are smaller than an upper bound *k*. In both cases, SPADIS selects *k* SNPs.

Some of the methods that we compare SPADIS to, such as SConES and Lasso, do not operate with a cardinality constraint directly. In order to satisfy the tight cardinality constraint, during parameter selection of these methods, we apply binary search over a range of sparsity parameter values that yields numbers close to *k*. For the rest of the parameters or all parameters in the case of maximum cardinality constraint (including sparsity parameter), we select them using two metrics separately: *stability*, denoted with (S) and found using the consistency index as described in Kuncheva (2007), and *regression performance*, denoted with (R), measured using Pearson’s squared correlation coefficient. The details on parameter selection for each method are provided in Supplementary Text 3.1.

Since we compare SPADIS with SConES in various settings, as a first step, we verify that we make use of SConES properly by replicating the results reported in Azencott *et al*. (2013) using their setting. Then, we compare SPADIS with SConES and other methods using another evaluation scheme.

*1) Replicating results of SConES:* Here, we use SConES’ setting explained in Azencott *et al*. (2013). First, using 10-fold cross validation, the desired objective function (i.e. stability for SConES(S), regression performance for SConES(R)) are measured for all parameters tested. The parameter values that maximize the desired objective are selected, and the final SNP set is determined with these parameters. Then, for evaluation, a ridge regression is performed on the complete dataset in a 10-fold cross validated setting using this SNP set and Pearson’s squared correlation coefficient is calculated for regression performance. Although this strategy is adopted by Azencott *et al*. (2013) due to the limited dataset size, it also implicates that the test data is used during the parameter selection step which might lead to memorization.

In order to reproduce the results, we apply tight cardinality constraint during parameter selection, targeted at the number of SNPs that are reported in the paper. We show that our replicated results are on par with the reported *R*^2^ and ratio of SNPs near candidate genes, respectively, indicating that we are able to replicate their results. These results are shown in Supplementary Figures 1 and 2, respectively. In addition, we run SConES(R), SConES(S) and SPADIS for the tight cardinality constraint of *k* = 500 using this setting. The corresponding results suggest that SPADIS performs better in regression performance in this setting —see Supplementary Figure 3.

**Fig. 1:**
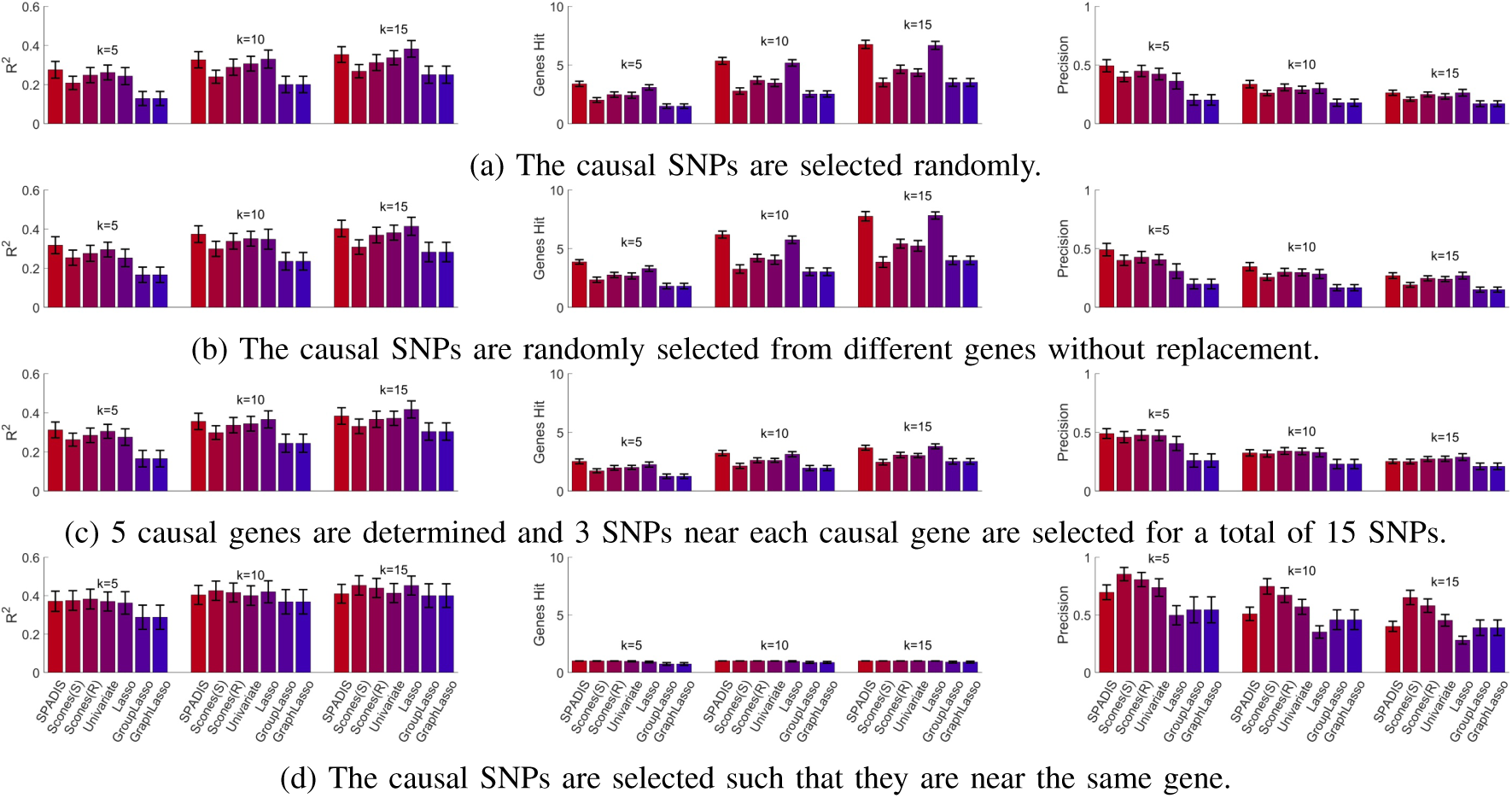
The simulation results of SPADIS, SConES(S), SConES(R), Univariate, Lasso, GroupLasso and GraphLasso for *k* = 5, *k* = 10 and *k* = 15. (a) Causal SNPs are picked randomly. (b) All causal SNPs are from different genes. (c) Causal SNPs are from 5 different genes (d) All causal SNPs are from the same gene. (Left) Pearson’s squared correlation coefficient, (Middle) Number of causal genes hit, (Right) Precision calculated for causal SNPs hit. Black bars indicate the 95% confidence intervals.

**Fig. 2:**
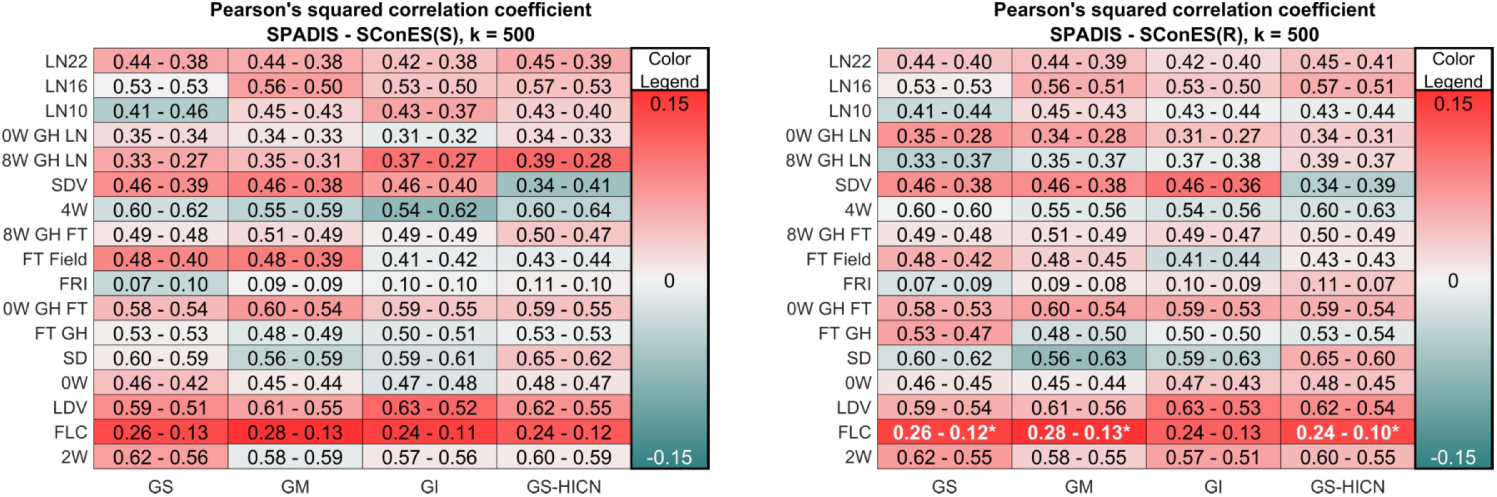
The regression performance comparisons of SPADIS with SConES(S) and SConES(R) on AT data for tight cardinality constraint of *k* = 500. The rows denote phenotypes and the columns denote networks. The numbers in each cell show Pearson’s squared correlation coefficients attained by SPADIS and SConES respectively. The background color encodes the difference in correlation coefficients. Red indicates SPADIS performs better than SConES while blue indicates otherwise. Differences that are found to be statistically significant are shown in bold, white font and marked with star (*).

**Fig. 3:**
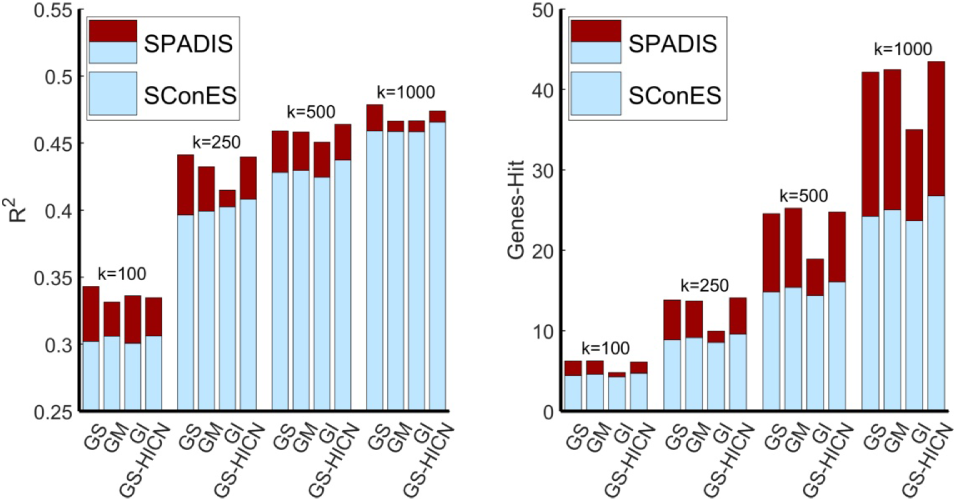
The improvement of SPADIS over SConES in terms of (left) Pearson’s squared correlation coefficient and (right) number of distinct candidate genes-hit for different tight cardinality constaints *k*. All values shown are averages over 17 phenotypes. Blue bar indicates the maximum of SConES(S) and SConES(R) for the corresponding network and *k* value. The red bar indicates the amount of improvement of SPADIS over SConES.

*2) Evaluation of SPADIS and compared methods:* In this study, we use nested cross-validation for evaluation. The outer 10-fold cross-validation splits the data into training and test data, and the inner 10-fold cross-validation selects the parameters using the training data only. For each fold in the outer cross-validation, a separate SNP set is selected and the test data is not seen by the algorithms. Unless otherwise stated, we use this setting in our experiments.

### E. Simulation Experiments

To assess the performance of the methods in a controlled manner, we conduct simulation experiments. We randomly choose 200 samples (out of 1307) in *AT* data. We select 500 random SNPs with MAF > 10% as follows: We first select 25 genes randomly. Then, we select 20 random SNPs near (< 20 kbp) each gene. In each experiment, we designate 15 SNPs to be causal and generate phenotypes using the regression model: **y** = **Xw** + *ϵ*, where 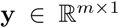 is the phenotype vector, 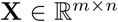 is the genotype matrix, 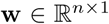 is the weight vector for each SNP, and *ϵ* is the error term. Both **w** and *ϵ* are normally distributed. We sample the weights of the causal SNPs from a standard normal distribution. We argue that in a real-life setting, there is no clear separation between causal and non-causal SNPs i.e. all SNPs play some part in explaining the phenotype at varying degrees. Hence, we sample the weights of the non-causal SNPs from a normal distribution with zero mean and 0.1 standard deviation instead of setting the standard deviations directly to zero. In our tests, we use the GS network as the SNP-SNP network.

We compare the methods under four different simulation settings: (a) the causal SNPs are randomly selected, (b) the causal SNPs are selected randomly such that they are near different genes, (c) 5 causal genes are determined and 3 SNPs near each causal gene are selected for a total of 15 SNPs, and (d) the causal SNPs are selected near a single random gene.

For each method, we adopt the tight cardinality constraint and test with *k* = 5, 10 and 15. For evaluation, we consider three metrics: (i) Precision as the ratio of the number of causal SNPs that are selected to the total number of SNPs selected, (ii) Number of causal genes hit (a gene is hit if a SNP near that gene is selected), (iii) Pearson’s squared correlation coefficient (*R*^2^). We perform 10-fold cross-validation 50 times and report averages over all folds. The 95% confidence interval for the means of the specified statistics are calculated assuming a t-distribution on the error.

In simulation settings (a), (b) and (c), SPADIS outperforms other methods when *k* is less than the number of causal SNPs —see Figure 1. When *k* is equal to the number of causal SNPs, Lasso catches up to SPADIS and they outperform all other methods. In setting (d) where the assumptions of SPADIS are violated, SPADIS underperforms compared to others in terms of Precision. Regardless, its regression performance is on a par with other methods. Note that, this is the setting where methods with graph connectivity assumption should perform well (casual SNPs are close). However, this scenario is not realistic since all associated SNPs are rarely that close to each other for complex traits.

Next, we check the number of causal genes hit (GenesHit) for all methods. In all simulation settings, we observe a correlation between GenesHit and *R*^2^ (i.e. methods that perform well in GenesHit perform well in *R*^2^ as well). We argue that high number of hit genes indicates high regression performance because when the selected SNPs fall into different genes, they are likely to contain complementary information and can explain the phenotype better. This constitutes the core idea of SPADIS.

### F. Phenotype Prediction Performance

*1) Experiments with Tight Cardinality Constraint:* First, we compare the regression performances of SConES(S), SConES(R) and SPADIS in *AT* data using the Pearson’s squared correlation coefficient (*R*^2^) by constraining them to select close to *k* SNPs (tight cardinality constraint). Here, we report results for *k* = 500 which we consider representative —see Figure 2. The results for *k* = 100, 250, and 1000 are provided in Supplementary Figures 4, 5 and 6, respectively.

**Fig. 4:**
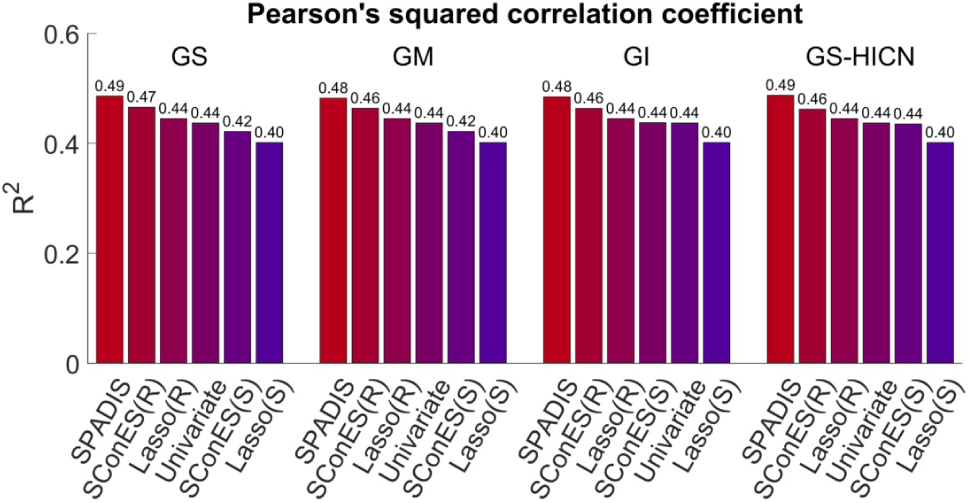
Regression performances of SPADIS, SConES(S), SConES(R), Univariate, Lasso(S) and Lasso(R) averaged over 17 AT phenotypes for maximum cardinality constraint of 1733. X-axis shows the compared methods and Y-axis shows the Pearson’s squared correlation coefficient (*R*^2^). For each network, methods are ordered in descending order of *R*^2^.

**Fig. 5:**
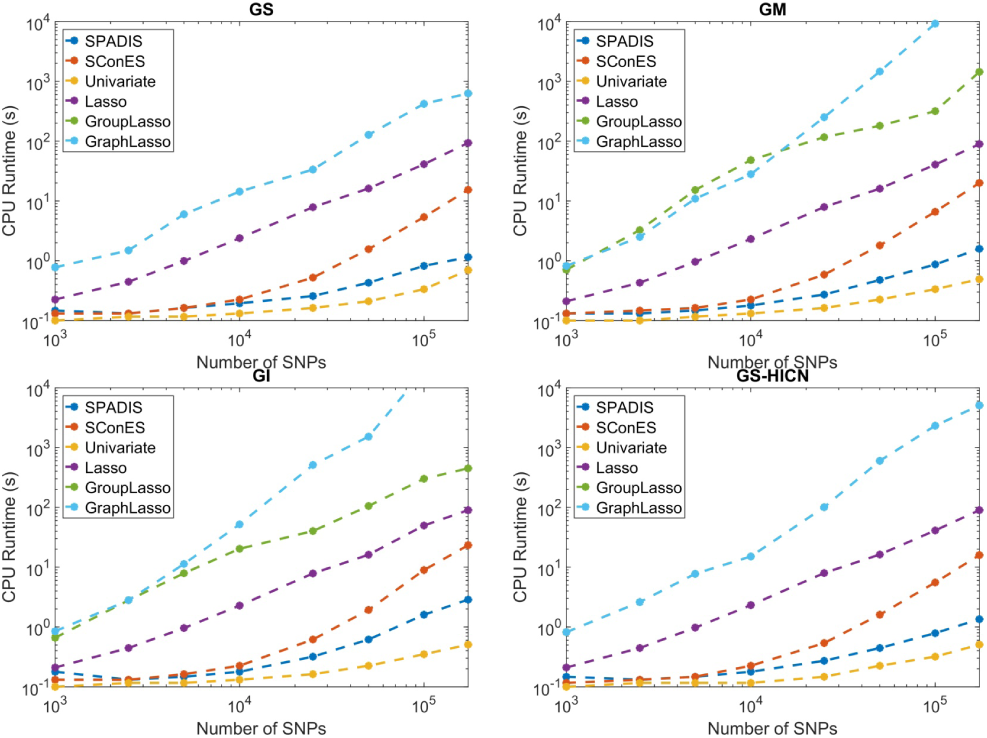
CPU time measurements of SPADIS, SConES, Univariate, Lasso, GroupLasso and GraphLasso from 1.000 to 173.219 SNPs on four networks: (Top left) GS, (Top right) GM, (Bottom left) GI and (Bottom right) GS-HICN. Note that, runtimes of GroupLasso and GraphLasso are the same for GS and GS-HICN networks by construction.

Out of 68 tests that is performed for *k* = 500 over 17 phenotypes using 4 different networks separately as input, SPADIS outperforms SConES(S) in 46 tests and SConES(R) in 47 tests. The improvement in *R*^2^ is up to 0.15 in a single phenotype and 0.03 on average. Overall, this corresponds to an improvement in 12 out of 17 phenotypes when averaged over all networks. Next, we test whether the differences in *R*^2^ are statistically significant (FDR adjusted p-value ≤ 0.05) using the method described in Hittner *et al*. (2003). The multiple hypothesis correction is conducted as in Yekutieli and Benjamini (1999). 3 results of SPADIS are found to be significantly better than SConES, whereas none of the results of SConES is found to be significantly better than SPADIS.

When averaged over all *k* values tested and all networks, SPADIS performs better than SConES in terms of Pearson’s squared correlation coefficient in 15 out of 17 phenotypes —see Supplementary Table 1. Moreover, SPADIS provides a consistent improvement in regression performance over SConES when averaged over all phenotypes. This improvement of SPADIS over SConES is summarized in Figure 3 for each network and *k* value tested. Note that, the improvement is particularly prevalent when *k* is smaller. On the other hand, we observe that average performance of both methods increase as the set size grows. Therefore, for a fair comparison, we believe that it is important to compare the methods when they select the same number of SNPs. That is why we perform the experiments with tight cardinality constraints.

**TABLE I:** Table shows statistics about the genes and biological processes hit by the selected SNPs sets by SPADIS, SConES(S), SConES(R), Univariate and Lasso. Tight cardinality constraint is applied for the following *k* values: *k* = 100, 250, 500 and 1000. The reported results are averages over all 17 phenotypes and 4 networks. The best result for each *k* is marked as bold.

| Metric | $k$ | SPADIS | SConES(S) | SConES(R) | Univariate | Lasso |
| --- | --- | --- | --- | --- | --- | --- |
| Genes-Hit | 100 | **5.9** | 4.4 | 4.5 | 3.8 | 5.5 |
|  | 250 | **12.9** | 8.7 | 9.0 | 7.6 | 10.9 |
|  | 500 | **23.4** | 14.3 | 15.0 | 13.8 | 18.3 |
|  | 1000 | **40.8** | 24.7 | 23.6 | 24.2 | 27.9 |
| GO-Hit | 100 | **151** | 114 | 117 | 137 | 144 |
|  | 250 | **306** | 230 | 236 | 266 | 280 |
|  | 500 | **491** | 373 | 382 | 424 | 441 |
|  | 1000 | **747** | 597 | 581 | 659 | 636 |
| Precision | 100 | 7.0% | **11.0%** | 10.9% | 8.6% | 7.3% |
|  | 250 | 6.3% | **9.4%** | 9.4% | 7.4% | 6.1% |
|  | 500 | 6.2% | 8.3% | **8.5%** | 6.9% | 5.9% |
|  | 1000 | 6.3% | 7.5% | **7.6%** | 6.7% | 5.8% |

*2) Experiments with Maximum Cardinality Constraint:* A more natural setting for SConES and other compared methods is to let them decide the number of SNPs based on their parameter search procedure. Hence, we perform a second set of experiments in which we allow methods to pick the SNP set size as long as the set sizes are bounded from above by 1733 i.e. 1% of the number of all SNPs as done in Azencott *et al*. (2013). Here, we compare SPADIS with SConES(S), SConES(R), Univariate, Lasso(S) and Lasso(R) on all phenotypes and all networks.

SPADIS is the best performing method in 8 out of 17 phenotypes on GS and GI networks and the best in 9 phenotypes on GM and GS-HICN networks (see Supplementary Figures 7–10). When regression performances (*R*^2^) are averaged over all phenotypes for each method, SPADIS outperforms all other methods on every network (see Figure 4). The next two best performing methods are SConES(R) and Lasso(R) respectively. Unsurprisingly, the methods that directly optimize or are tuned based on *R*^2^ are better in regression than their stability optimizing versions on average.

Next, we check whether the differences between SPADIS and other methods are statistically significant. Out of 68 experiments of SPADIS (17 phenotypes × 4 networks) SPADIS is found to be significantly better than (i) SConES(R) in 2 experiments, (ii) Lasso(R) in 6 experiments, (iii) SConES(S) in 14 experiments, (iv) Univariate in 17 experiments, and finally, (v) Lasso(S) in 28 experiments. In none of the experiments, SPADIS is found to be significantly worse than its counterparts. See Supplementary Figures 11–13 for the corresponding results.

### G. Diverse Selection of SNPs

The goal of SPADIS is to select a diverse set of SNPs over the SNP-SNP network. We hypothesize that SNPs selected with SPADIS overlap with more diverse biological processes and that the prediction performance is reinforced by this effect. Here, we investigate whether this hypothesis is supported by empirical values on the 17 flowering time phenotypes of *AT*. To this end, we utilize three metrics: (1) Genes-Hit, (2) GO-Hit, and (3) Precision, which are explained in the following subsections. Since the performance with respect to these metrics typically depends on the number of SNPs selected, we apply tight cardinality constraint and report the results for *k* = 100, 250, 500 and 1000.

*1) Evaluation with Genes-Hit metric:* First, we compare the average number of candidate genes hit by each method (out of 165 candidate genes related with flowering time). A gene is considered *hit* if the method selects a SNP *near* the gene (≤ 20 kbp). SPADIS hits 7%-46% more distinct candidate genes compared to the next best performing method on average, over different cardinality constraints —see Table I. This is an indication that SPADIS realizes one of its goals which is to spatially *cover* the network and genome.

*2) Evaluation with GO-Hit metric:* Here, we check how many distinct GO biological processes are hit by the SNPs selected by each method. A process is considered hit if the method chooses a SNP near a gene which is annotated with that biological term.

As shown in Table I, SNPs discovered by SPADIS covers 151, 306, 491 and 747 GO-terms on average for *k* = 100, 250, 500 and 1000 respectively. This is an increase of 5% to 17% compared to the next best performing method, over different cardinality constraints. It supports our intuition that SPADIS discovers SNPs that are related to diverse processes.

*3) Evaluation with Precision metric:* Finally, for the sake of completeness, we compare SPADIS and other methods with respect to the ratio of the number of selected SNPs that are near a candidate gene to the total number of selected SNPs, as done in Azencott *et al*. (2013). This metric measures the *precision* of the selected SNPs, hence we denote it as such. As shown in Table I, SPADIS consistently underperforms in this metric. Nevertheless, we argue that it is not a good measure of how well the methods perform. Precision considers all SNPs near a candidate gene as true positives. Consider the following extreme case: a method that selects solely a set of SNPs near a single candidate gene can achieve a precision of 1. Hence, precision indirectly rewards the selection of SNPs that fall into a smaller number of genes. On the other hand, the diversification of SNPs in terms of genes and biological processes help explain the phenotype better. This metric is in clear contrast with the number of genes hit and the number of biological processes hit.

### H. Contribution of the Hi-C Data

We evaluate the information leveraged by using the Hi-C data via comparing the regression performances obtained when using GS-HICN compared to using other networks (GS, GM, GI). Tests are performed for all 17 phenotypes with SPADIS, SConES(S) and SConES(R). We compared the methods over five experiments: four experiments with tight cardinality constraint applied for *k* = 100, 250, 500 and 1000, and one experiment with maximum cardinality constraint applied for *k* = 1733. As shown in Table II, Hi-C data provides improvements in regression performance on average: 1.4% higher than GS and GM and 1.9% higher than GI. Moreover, the improvement can be considered consistent since GS-HICN performs better than other networks on average in 4 out of 5 experiments. Moreover, GS-HICN hits 3.0% to 6.6% more genes and 2.7% to 21.9% more biological processes compared to other networks, on average — see Supplementary Tables 2–5. For comparisons of GS-HICN with other networks per individual phenotype in terms of regression performance, see Supplementary Figures 14–17.

**TABLE II:** Table shows the average Pearson’s squared correlation coefficient obtained for all networks and experiments that are tested. The results are averaged over all 17 phenotypes and all methods (SPADIS, SConES(S) and SConES(R)). The best result for each experiment is marked as bold.

| Experiment<br>Constraint | k | Network |  |  |  |
| --- | --- | --- | --- | --- | --- |
|  |  | GS | GM | GI | GS-HICN |
| Tight | 100 | 0.310 | 0.311 | 0.309 | **0.314** |
| Tight | 250 | 0.403 | 0.406 | 0.398 | **0.415** |
| Tight | 500 | 0.438 | 0.438 | 0.432 | **0.445** |
| Tight | 1000 | 0.461 | 0.461 | 0.459 | **0.467** |
| Maximum | 1733 | 0.457 | 0.456 | **0.462** | 0.461 |
| Average |  | 0.414 | 0.414 | 0.412 | **0.420** |

### I. Time Performance

We report the CPU runtime of all methods, across a range of number of SNPs (from 1000 to 173 219) and all four networks. The measurements are taken on a single dedicated core of Intel i7–6700HQ processor. The runtime tests are conducted for one cross-validation fold with preset parameters on a single phenotype FT Field, which has the most number of samples available (*m* = 180).

We consider a method to time-out if it takes more than 10^3^ seconds for a single run because the runtime of the complete test (10 folds with parameter selection) would take more than 1 CPU week (10^3^ seconds x 10 evaluation folds x 10 training folds x at least 7 parameters).

Results show that SPADIS is more efficient than all other methods except the Univariate (baseline) method —see Figure 5. GroupLasso and GraphLasso do not scale to SNP selection problem in GWAS. For this reason, they are not included in the experiments performed on *AT* data.

## IV. Discussion

SPADIS seeks for a subset of SNPs on a network derived from biological knowledge, such that the selected SNP set is associated with the phenotype. Even though there are other network based methods for tackling the same problem, they rest on the assumption that causal SNPs tend to be connected on the network. Thus, they incorporate constraints that favor the connectivity of selected SNPs. However, we argue that selecting connected SNPs together might not provide additional predictive power as they can be in haplotype blocks and bring redundant information. Moreover, a method that highlights different parts of the network could be useful because it can potentially recover different biological processes: SNPs affecting diverse biological processes would be complementary and explain the phenotype better. To address these issues, we propose a new formulation: As opposed to enforcing graph connectivity over the set of selected features, we set out to discover SNPs that are far apart in terms of their location on the genome, which translate into diversity in function. To the best of our knowledge, none of the current approaches operate with this principle. Our results indicate that selecting SNPs remotely located on the network indeed hit genes that are related to a larger number of distinct biological processes. This property can help in gaining more biological insights into the genetic basis of the complex traits and diseases.

The technical contribution of this paper involves formulating this principle through a submodular function. We empirically show that SPADIS can recover SNPs known to be associated with the phenotype and the optimization is efficient. Another alternative would be to formulate an optimization function that directly rewards the number of distinct process hits. However, given the incomplete knowledge of the process annotations, this could lead to literature bias. Therefore, we refrain from incorporating such a term directly in the model, instead, we let the diversity on the 2D and 3D locations lead the diverse selection.

In our experiments, to score each SNPs relevance to the phenotype, we use sequence kernel association test (SKAT) based on its success and for drawing a fair comparison to the literature. There are other alternatives such as Pearson’s correlation coefficient, or maximal information coefficient (Reshef *et al*., 2011), which can easily be used with SPADIS as long as the computed scores are non-negative or are transformed to a non-negative range.

For the first time, we investigate the utility of Hi-C data for selecting a SNP set. Our results show that Hi-C data consistently provides slight improvements in regression performance. We think it is a promising source of information for SNP association. We currently limit the use of data to intra-chromosomal contacts due to much better higher resolution compared to inter-chromosomal contact maps (2 kbp vs. 20 kbp). We also discard contacts that fall outside of the significance range. These choices are likely to over-constrain the method, and further research is needed to fully utilize such information, which we leave as future work.

In this article, we introduce SPADIS and benchmark its performance on *AT* genotype and phenotypes. Alternatively, SPADIS can be used for discovering associated SNP sets for complex genetic disorders as well. For instance in autism, research efforts have mostly focused on identifying risk genes through whole exome sequencing studies (De Rubeis *et al*., 2014; Iossifov *et al*., 2014). However, close to 90% of the point mutations fall outside of the coding regions (Hindorff *et al*., 2009). Discovering a set of non-coding risk mutations will certainly help to uncover the genetic architecture. Recently, a large-scale effort to collect GWAS data of autism families along with clinical information of patients is reported (Yuen *et al*., 2017). Hence, in future work, we plan to apply SPADIS on autism, which should help explain the heterogeneity in wide spectrum of phenotypes.

## Acknowledgements

We thank Chlo-Agathe Azencott and Dominik Grimm for their help on running SConES, Mehmet Koyutrk and Utku Norman for their feedback on SPADIS. We thank TUBITAK for supporting this research via Career Grant #116E148 to AEC.

## Footnotes

1 ftp://ftp.arabidopsis.org/home/tair/Proteins/

