## Supplementary Materials for "SPADIS: An Algorithm for Selecting Predictive and Diverse SNPs in GWAS" (from another version of this preprint: version 2, 2018-03-26)

#### 1 Supplementary Figures

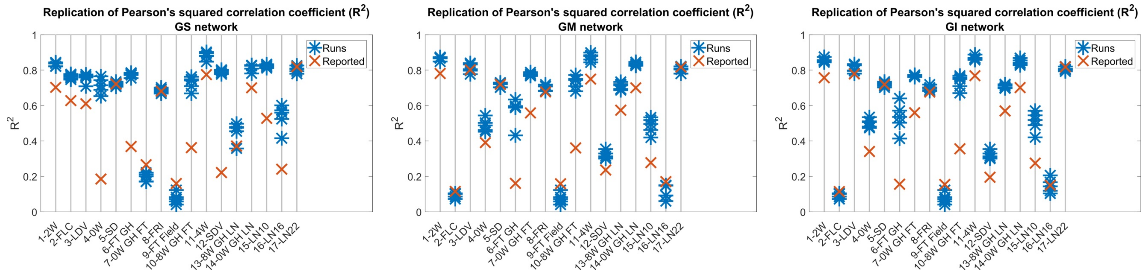


Figure 1: Figure shows the replication of the Pearson's squared correlation coefficients ($R^2$) results of Azencott et. al, 2013 successfully. We do not have the actual folds used in the paper. Using binary search, we adjust the parameters so that SConES selects the same the number of SNPs used in the paper to produce the reported $R^2$ value. Each sub-figure represents the underlying network: GS (left), GM (center), and GI (right). x and y axis show phenotypes and $R^2$ values for each phenotype respectively. Cross signs indicate the value reported in Azencott et. al, 2013, for that phenotype. Blue stars show the results we obtained for 5 different 10-fold cross validation splits. We show that the reported values can be reproduced in Experimental Setting 1 (ES1). We could not reproduce these results in ES2, where we perform nested 10-fold cross-validation.

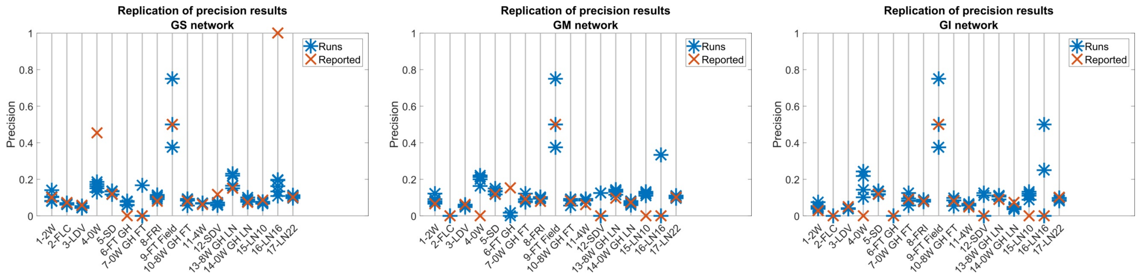


Figure 2: Figure shows the replication of the precision results of Azencott et. al, 2013. Precision is defined as the ratio of the number of SNPs selected that are near candidate genes and total number of SNPs selected. We do not have the actual folds used in the paper. Using binary search, we adjust the parameters so that SConES selects the same the number of SNPs used in the paper to produce the reported precision value. Each sub-figure represents the underlying network: GS (left), GM (center), and GI (right). x and y axis show phenotypes and precision for each phenotype respectively. Cross signs indicate the reported precision value in Azencott et. al, 2013, for that phenotype. Blue stars show the results we obtained for 5 different 10-fold cross validation splits. We show that the reported values can be reproduced in Experimental Setting 1 (ES1) as the reported values can be achieved in our runs. We could not reproduce these results in ES2.

**Pearson's squared correlation coefficient**
**SPADIS - SConES(S), k = 100**

| | GS | GM | GI | GS-HICN |
|---|---|---|---|---|
| LN22 | 0.33 - 0.31 | 0.36 - 0.29 | 0.32 - 0.31 | 0.33 - 0.30 |
| LN16 | 0.46 - 0.42 | 0.44 - 0.43 | 0.44 - 0.45 | 0.40 - 0.47 |
| LN10 | 0.28 - 0.21 | 0.28 - 0.23 | 0.30 - 0.26 | 0.29 - 0.29 |
| 0W GH LN | 0.28 - 0.20 | 0.29 - 0.23 | 0.33 - 0.17 | 0.20 - 0.18 |
| 8W GH LN | 0.29 - 0.19 | 0.31 - 0.21 | 0.29 - 0.22 | 0.22 - 0.16 |
| SDV | 0.33 - 0.16 | **0.34 - 0.15*** | 0.34 - 0.18 | 0.21 - 0.21 |
| 4W | 0.42 - 0.43 | 0.34 - 0.44 | 0.37 - 0.36 | 0.53 - 0.44 |
| 8W GH FT | 0.38 - 0.34 | 0.42 - 0.32 | 0.41 - 0.35 | 0.41 - 0.32 |
| FT Field | 0.29 - 0.23 | 0.31 - 0.25 | 0.32 - 0.21 | 0.30 - 0.27 |
| FRI | 0.08 - 0.07 | 0.07 - 0.06 | 0.11 - 0.11 | 0.13 - 0.07 |
| 0W GH FT | 0.34 - 0.33 | 0.35 - 0.32 | 0.38 - 0.33 | 0.37 - 0.35 |
| FT GH | 0.48 - 0.37 | 0.41 - 0.39 | 0.44 - 0.41 | 0.44 - 0.43 |
| SD | 0.58 - 0.55 | 0.55 - 0.57 | 0.53 - 0.56 | 0.55 - 0.57 |
| 0W | 0.34 - 0.25 | 0.27 - 0.23 | 0.20 - 0.27 | 0.30 - 0.27 |
| LDV | 0.42 - 0.45 | 0.46 - 0.52 | 0.46 - 0.43 | 0.42 - 0.40 |
| FLC | 0.13 - 0.07 | 0.12 - 0.06 | 0.12 - 0.06 | 0.22 - 0.08 |
| 2W | 0.39 - 0.31 | 0.32 - 0.33 | 0.37 - 0.27 | 0.40 - 0.34 |

Color Legend: 0.19 / 0 / -0.19

**Pearson's squared correlation coefficient**
**SPADIS - SConES(R), k = 100**

| | GS | GM | GI | GS-HICN |
|---|---|---|---|---|
| LN22 | 0.33 - 0.37 | 0.36 - 0.37 | 0.32 - 0.36 | 0.33 - 0.37 |
| LN16 | 0.46 - 0.44 | 0.44 - 0.44 | 0.44 - 0.44 | 0.40 - 0.43 |
| LN10 | 0.28 - 0.20 | 0.28 - 0.21 | 0.30 - 0.20 | 0.29 - 0.22 |
| 0W GH LN | 0.28 - 0.28 | 0.29 - 0.29 | 0.33 - 0.24 | 0.20 - 0.28 |
| 8W GH LN | 0.29 - 0.28 | 0.31 - 0.27 | 0.29 - 0.28 | 0.22 - 0.26 |
| SDV | **0.33 - 0.17*** | **0.34 - 0.17*** | 0.34 - 0.20 | 0.21 - 0.16 |
| 4W | 0.42 - 0.39 | 0.34 - 0.41 | 0.37 - 0.41 | 0.53 - 0.40 |
| 8W GH FT | 0.38 - 0.37 | 0.42 - 0.37 | 0.41 - 0.36 | 0.41 - 0.35 |
| FT Field | 0.29 - 0.23 | 0.31 - 0.23 | 0.32 - 0.25 | 0.30 - 0.26 |
| FRI | 0.08 - 0.08 | 0.07 - 0.10 | 0.11 - 0.10 | 0.13 - 0.10 |
| 0W GH FT | 0.34 - 0.35 | 0.35 - 0.35 | 0.38 - 0.35 | 0.37 - 0.34 |
| FT GH | 0.48 - 0.39 | 0.41 - 0.39 | 0.44 - 0.35 | 0.44 - 0.42 |
| SD | 0.58 - 0.58 | 0.55 - 0.58 | 0.53 - 0.55 | 0.52 - 0.56 |
| 0W | 0.34 - 0.21 | 0.27 - 0.20 | 0.20 - 0.22 | 0.30 - 0.23 |
| LDV | 0.42 - 0.44 | 0.46 - 0.46 | 0.46 - 0.44 | 0.42 - 0.44 |
| FLC | 0.13 - 0.06 | 0.12 - 0.06 | 0.12 - 0.06 | 0.22 - 0.10 |
| 2W | 0.39 - 0.30 | 0.32 - 0.31 | 0.37 - 0.29 | 0.40 - 0.28 |

Color Legend: 0.17 / 0 / -0.17

Figure 3: The regression performances of SPADIS, SConES(R) and SConES(S) on AT data for PSS1 and $k = 100$. Rows represent different phenotypes and columns represent different networks. The numbers in each cell show Pearson's squared correlation coefficients achieved by SPADIS and SConES, respectively (separated with a dash). The tone of background color reflects the difference in correlation coefficients between SPADIS and SConES. Red indicates SPADIS performs better than SConES while blue indicates otherwise. Differences that are found to be statistically significant are shown in bold and white and marked with star (*).

**Pearson's squared correlation coefficient**
**SPADIS - SConES(S), k = 250**

| | GS | GM | GI | GS-HICN |
|---|---|---|---|---|
| LN22 | 0.42 - 0.37 | 0.40 - 0.37 | 0.37 - 0.36 | 0.40 - 0.37 |
| LN16 | 0.52 - 0.47 | 0.50 - 0.50 | 0.47 - 0.48 | 0.53 - 0.52 |
| LN10 | 0.39 - 0.37 | 0.41 - 0.36 | 0.37 - 0.36 | 0.49 - 0.39 |
| 0W GH LN | 0.37 - 0.28 | 0.38 - 0.30 | 0.31 - 0.24 | 0.31 - 0.30 |
| 8W GH LN | 0.35 - 0.24 | 0.35 - 0.27 | 0.35 - 0.27 | 0.35 - 0.21 |
| SDV | **0.45 - 0.28*** | **0.49 - 0.30*** | **0.46 - 0.28*** | 0.36 - 0.35 |
| 4W | 0.58 - 0.49 | 0.54 - 0.53 | 0.59 - 0.49 | 0.55 - 0.63 |
| 8W GH FT | 0.49 - 0.42 | 0.45 - 0.41 | 0.45 - 0.42 | 0.49 - 0.40 |
| FT Field | 0.45 - 0.35 | 0.43 - 0.40 | 0.40 - 0.37 | 0.48 - 0.37 |
| FRI | 0.08 - 0.08 | 0.06 - 0.08 | 0.13 - 0.10 | 0.09 - 0.09 |
| 0W GH FT | 0.54 - 0.46 | 0.56 - 0.45 | 0.56 - 0.45 | 0.59 - 0.46 |
| FT GH | 0.52 - 0.49 | 0.52 - 0.55 | 0.47 - 0.54 | 0.48 - 0.56 |
| SD | 0.59 - 0.61 | 0.58 - 0.57 | 0.55 - 0.59 | 0.61 - 0.63 |
| 0W | 0.42 - 0.36 | 0.33 - 0.37 | 0.35 - 0.40 | 0.43 - 0.39 |
| LDV | 0.54 - 0.47 | 0.60 - 0.51 | 0.52 - 0.48 | 0.58 - 0.49 |
| FLC | 0.23 - 0.11 | 0.24 - 0.12 | 0.17 - 0.10 | 0.22 - 0.12 |
| 2W | 0.56 - 0.46 | 0.52 - 0.48 | 0.53 - 0.47 | 0.52 - 0.47 |

Color Legend: 0.18 / 0 / -0.18

**Pearson's squared correlation coefficient**
**SPADIS - SConES(R), k = 250**

| | GS | GM | GI | GS-HICN |
|---|---|---|---|---|
| LN22 | 0.42 - 0.41 | 0.40 - 0.41 | 0.37 - 0.42 | 0.40 - 0.41 |
| LN16 | 0.52 - 0.53 | 0.50 - 0.53 | 0.47 - 0.54 | 0.53 - 0.53 |
| LN10 | 0.39 - 0.40 | 0.41 - 0.42 | 0.37 - 0.42 | 0.49 - 0.42 |
| 0W GH LN | 0.37 - 0.32 | 0.38 - 0.34 | 0.31 - 0.33 | 0.31 - 0.33 |
| 8W GH LN | 0.35 - 0.33 | 0.35 - 0.34 | 0.35 - 0.33 | 0.35 - 0.34 |
| SDV | 0.45 - 0.31 | **0.49 - 0.32*** | 0.46 - 0.32 | 0.36 - 0.32 |
| 4W | 0.58 - 0.55 | 0.54 - 0.53 | 0.59 - 0.53 | 0.55 - 0.59 |
| 8W GH FT | 0.49 - 0.42 | 0.45 - 0.42 | 0.45 - 0.43 | 0.49 - 0.43 |
| FT Field | 0.45 - 0.36 | 0.43 - 0.35 | 0.40 - 0.37 | 0.48 - 0.37 |
| FRI | 0.08 - 0.08 | 0.06 - 0.09 | 0.13 - 0.10 | 0.09 - 0.08 |
| 0W GH FT | 0.54 - 0.43 | 0.56 - 0.44 | 0.56 - 0.44 | **0.59 - 0.43*** |
| FT GH | 0.52 - 0.47 | 0.52 - 0.48 | 0.47 - 0.48 | 0.48 - 0.49 |
| SD | 0.59 - 0.64 | 0.58 - 0.63 | 0.55 - 0.62 | 0.61 - 0.65 |
| 0W | 0.42 - 0.36 | 0.33 - 0.37 | 0.35 - 0.37 | 0.43 - 0.40 |
| LDV | 0.54 - 0.53 | 0.60 - 0.53 | 0.52 - 0.53 | 0.58 - 0.54 |
| FLC | 0.23 - 0.12 | 0.24 - 0.12 | 0.17 - 0.13 | 0.22 - 0.13 |
| 2W | 0.56 - 0.46 | 0.52 - 0.47 | 0.53 - 0.48 | 0.52 - 0.47 |

Color Legend: 0.17 / 0 / -0.17

Figure 4: The regression performances of SPADIS, SConES(R) and SConES(S) on AT data for PSS1 and $k = 250$. Rows represent different phenotypes and columns represent different networks. The numbers in each cell show Pearson's squared correlation coefficients achieved by SPADIS and SConES, respectively (separated with a dash). The tone of background color reflects the difference in correlation coefficients between SPADIS and SConES. Red indicates SPADIS performs better than SConES while blue indicates otherwise. Differences that are found to be statistically significant are shown in bold and white and marked with star (*).

**Pearson's squared correlation coefficient**
**SPADIS - SConES(S), k = 1000**

| | GS | GM | GI | GS-HICN |
|---|---|---|---|---|
| LN22 | 0.48 - 0.41 | 0.42 - 0.40 | 0.43 - 0.40 | 0.47 - 0.42 |
| LN16 | 0.55 - 0.52 | 0.52 - 0.54 | 0.54 - 0.53 | 0.57 - 0.55 |
| LN10 | 0.42 - 0.42 | 0.45 - 0.42 | 0.46 - 0.40 | 0.49 - 0.44 |
| 0W GH LN | 0.45 - 0.35 | 0.37 - 0.40 | 0.40 - 0.38 | 0.37 - 0.38 |
| 8W GH LN | 0.34 - 0.34 | 0.34 - 0.33 | 0.34 - 0.33 | 0.36 - 0.35 |
| SDV | 0.47 - 0.40 | 0.42 - 0.40 | 0.44 - 0.41 | 0.36 - 0.40 |
| 4W | 0.60 - 0.63 | 0.59 - 0.64 | 0.61 - 0.60 | 0.60 - 0.66 |
| 8W GH FT | 0.46 - 0.49 | 0.45 - 0.48 | 0.46 - 0.50 | 0.45 - 0.47 |
| FT Field | 0.48 - 0.47 | 0.46 - 0.47 | 0.46 - 0.48 | 0.47 - 0.46 |
| FRI | 0.12 - 0.10 | 0.09 - 0.11 | 0.09 - 0.11 | 0.11 - 0.12 |
| 0W GH FT | 0.64 - 0.58 | 0.66 - 0.59 | 0.58 - 0.60 | 0.61 - 0.61 |
| FT GH | 0.55 - 0.53 | 0.53 - 0.53 | 0.56 - 0.54 | 0.56 - 0.54 |
| SD | 0.61 - 0.61 | 0.60 - 0.62 | 0.58 - 0.61 | 0.64 - 0.63 |
| 0W | 0.48 - 0.43 | 0.46 - 0.49 | 0.46 - 0.45 | 0.50 - 0.49 |
| LDV | 0.57 - 0.57 | 0.65 - 0.58 | 0.62 - 0.54 | 0.64 - 0.61 |
| FLC | 0.28 - 0.15 | 0.29 - 0.18 | 0.30 - 0.18 | 0.22 - 0.16 |
| 2W | 0.64 - 0.60 | 0.63 - 0.60 | 0.61 - 0.61 | 0.63 - 0.62 |

Color Legend: 0.12 / 0 / -0.12

**Pearson's squared correlation coefficient**
**SPADIS - SConES(R), k = 1000**

| | GS | GM | GI | GS-HICN |
|---|---|---|---|---|
| LN22 | 0.48 - 0.41 | 0.42 - 0.41 | 0.43 - 0.41 | 0.47 - 0.41 |
| LN16 | 0.55 - 0.54 | 0.52 - 0.51 | 0.54 - 0.51 | 0.57 - 0.55 |
| LN10 | 0.42 - 0.48 | 0.45 - 0.46 | 0.46 - 0.45 | 0.49 - 0.49 |
| 0W GH LN | 0.45 - 0.36 | 0.37 - 0.36 | 0.40 - 0.37 | 0.37 - 0.36 |
| 8W GH LN | 0.34 - 0.35 | 0.34 - 0.34 | 0.34 - 0.35 | 0.36 - 0.33 |
| SDV | 0.47 - 0.44 | 0.42 - 0.44 | 0.44 - 0.43 | 0.36 - 0.43 |
| 4W | 0.60 - 0.61 | 0.59 - 0.61 | 0.61 - 0.61 | 0.60 - 0.61 |
| 8W GH FT | 0.46 - 0.47 | 0.45 - 0.47 | 0.46 - 0.47 | 0.45 - 0.47 |
| FT Field | 0.48 - 0.47 | 0.46 - 0.48 | 0.46 - 0.48 | 0.47 - 0.48 |
| FRI | 0.12 - 0.10 | 0.09 - 0.11 | 0.09 - 0.09 | 0.11 - 0.09 |
| 0W GH FT | 0.64 - 0.60 | 0.66 - 0.60 | 0.58 - 0.60 | 0.61 - 0.60 |
| FT GH | 0.55 - 0.55 | 0.53 - 0.55 | 0.56 - 0.57 | 0.56 - 0.56 |
| SD | 0.61 - 0.62 | 0.60 - 0.62 | 0.58 - 0.62 | 0.64 - 0.62 |
| 0W | 0.48 - 0.45 | 0.46 - 0.48 | 0.46 - 0.46 | 0.50 - 0.45 |
| LDV | 0.57 - 0.58 | 0.65 - 0.56 | 0.62 - 0.56 | 0.64 - 0.59 |
| FLC | 0.28 - 0.17 | 0.29 - 0.19 | 0.30 - 0.20 | 0.22 - 0.16 |
| 2W | 0.64 - 0.61 | 0.63 - 0.61 | 0.61 - 0.60 | 0.63 - 0.62 |

Color Legend: 0.11 / 0 / -0.11

Figure 5: The regression performances of SPADIS, SConES(R) and SConES(S) on AT data for PSS1 and $k = 1000$. Rows represent different phenotypes and columns represent different networks. The numbers in each cell show Pearson's squared correlation coefficients achieved by SPADIS and SConES, respectively (separated with a dash). The tone of background color reflects the difference in correlation coefficients between SPADIS and SConES. Red indicates SPADIS performs better than SConES while blue indicates otherwise. Differences that are found to be statistically significant are shown in bold and white and marked with star (*).

**Pearson's squared correlation coefficient**
**SPADIS - SConES(S), k = 500**

| | GS | GM | GI | GS-HICN |
|---|---|---|---|---|
| LN22 | **0.84 - 0.75*** | **0.83 - 0.76*** | 0.80 - 0.76 | 0.81 - 0.76 |
| LN16 | **0.84 - 0.77*** | 0.83 - 0.77 | 0.83 - 0.78 | 0.82 - 0.79 |
| LN10 | 0.80 - 0.74 | 0.79 - 0.73 | 0.80 - 0.74 | 0.81 - 0.75 |
| 0W GH LN | 0.84 - 0.77 | **0.84 - 0.77*** | 0.85 - 0.84 | 0.80 - 0.81 |
| 8W GH LN | **0.80 - 0.66*** | **0.79 - 0.67*** | **0.79 - 0.67*** | **0.78 - 0.66*** |
| SDV | **0.87 - 0.80*** | 0.86 - 0.81 | 0.83 - 0.80 | 0.85 - 0.81 |
| 4W | 0.91 - 0.90 | 0.92 - 0.89 | 0.91 - 0.87 | 0.91 - 0.89 |
| 8W GH FT | **0.83 - 0.78*** | **0.83 - 0.78*** | **0.83 - 0.78*** | **0.82 - 0.78*** |
| FT Field | **0.82 - 0.75*** | **0.82 - 0.75*** | 0.80 - 0.75 | **0.79 - 0.72*** |
| FRI | 0.71 - 0.69 | 0.73 - 0.70 | 0.71 - 0.71 | 0.68 - 0.68 |
| 0W GH FT | **0.87 - 0.79*** | **0.88 - 0.78*** | **0.86 - 0.77*** | 0.84 - 0.79 |
| FT GH | **0.84 - 0.74*** | 0.82 - 0.79 | 0.84 - 0.80 | **0.85 - 0.78*** |
| SD | **0.86 - 0.81*** | **0.90 - 0.81*** | 0.85 - 0.81 | **0.90 - 0.81*** |
| 0W | **0.87 - 0.72*** | **0.87 - 0.79*** | **0.88 - 0.79*** | 0.86 - 0.82 |
| LDV | 0.80 - 0.77 | **0.81 - 0.74*** | **0.82 - 0.72*** | **0.83 - 0.76*** |
| FLC | **0.79 - 0.62*** | **0.81 - 0.70*** | **0.81 - 0.69*** | **0.75 - 0.65*** |
| 2W | 0.86 - 0.82 | **0.88 - 0.83*** | 0.86 - 0.82 | 0.87 - 0.84 |

Color Legend: 0.17, 0, -0.17

**Pearson's squared correlation coefficient**
**SPADIS - SConES(R), k = 500**

| | GS | GM | GI | GS-HICN |
|---|---|---|---|---|
| LN22 | **0.84 - 0.79*** | 0.83 - 0.79 | 0.80 - 0.79 | 0.81 - 0.79 |
| LN16 | **0.84 - 0.78*** | 0.83 - 0.78 | **0.83 - 0.78*** | 0.82 - 0.78 |
| LN10 | 0.80 - 0.75 | 0.79 - 0.75 | 0.80 - 0.75 | 0.81 - 0.77 |
| 0W GH LN | **0.84 - 0.74*** | **0.84 - 0.74*** | 0.85 - 0.80 | 0.80 - 0.82 |
| 8W GH LN | **0.80 - 0.73*** | **0.79 - 0.73*** | **0.79 - 0.73*** | **0.78 - 0.73*** |
| SDV | **0.87 - 0.80*** | **0.86 - 0.80*** | 0.83 - 0.80 | 0.85 - 0.80 |
| 4W | 0.91 - 0.90 | 0.92 - 0.91 | 0.91 - 0.90 | 0.91 - 0.90 |
| 8W GH FT | **0.83 - 0.79*** | **0.83 - 0.79*** | **0.83 - 0.79*** | 0.82 - 0.79 |
| FT Field | 0.82 - 0.77 | **0.82 - 0.76*** | 0.80 - 0.76 | 0.79 - 0.77 |
| FRI | 0.71 - 0.68 | 0.73 - 0.68 | 0.71 - 0.69 | 0.68 - 0.68 |
| 0W GH FT | **0.87 - 0.79*** | **0.88 - 0.80*** | **0.86 - 0.80*** | 0.84 - 0.80 |
| FT GH | 0.84 - 0.81 | 0.82 - 0.81 | 0.84 - 0.79 | 0.85 - 0.81 |
| SD | **0.86 - 0.81*** | **0.90 - 0.81*** | **0.85 - 0.81*** | **0.90 - 0.81*** |
| 0W | **0.87 - 0.75*** | **0.87 - 0.75*** | **0.88 - 0.78*** | 0.86 - 0.81 |
| LDV | **0.80 - 0.73*** | **0.81 - 0.73*** | **0.82 - 0.74*** | **0.83 - 0.76*** |
| FLC | **0.79 - 0.72*** | **0.81 - 0.72*** | **0.81 - 0.70*** | **0.75 - 0.65*** |
| 2W | 0.86 - 0.86 | 0.88 - 0.86 | 0.86 - 0.86 | 0.87 - 0.85 |

Color Legend: 0.12, 0, -0.12

Figure 6: The regression performances of SPADIS, SConES(R) and SConES(S) on AT data for PSS1 and $k = 500$, when the Experimental Setting 1 (ES1) is used as described in the main text. The rows denote phenotypes and columns denote networks. The numbers in each cell show Pearson's squared correlation coefficients ($R^2$) attained by SPADIS and SConES, respectively. The background color reflects the difference in Pearson's squared correlation coefficients between SPADIS and SConES. Red indicates SPADIS performs better than SConES while blue indicates otherwise. Differences that are found to be statistically significant are shown in bold and white and marked with star (*). Note that $R^2$ values and SConES' improvement are much larger compared to ES2 as the test data leaks into the feature selection procedure.

**Pearson's squared correlation coefficient, GM**

| | SPADIS | SConES(S) | SConES(R) | Univariate | Lasso(R) | Lasso(S) |
|---|---|---|---|---|---|---|
| LN22 | **0.45 (1733)** | 0.35 (1398) | 0.41 (1233) | 0.34 (1733) | 0.38 (414) | 0.44 (826) |
| LN16 | **0.58 (1733)** | 0.57 (1466) | 0.52 (1362) | 0.53 (1733) | 0.57 (1248) | 0.41 (814) |
| LN10 | 0.45 (1733) | 0.43 (1346) | 0.48 (1288) | 0.42 (1733) | **0.49 (347)** | 0.35 (510) |
| 0W GH LN | **0.39 (1733)** | 0.29 (1049) | 0.36 (1050) | 0.32 (360) | **0.39 (755)** | 0.35 (1274) |
| 8W GH LN | **0.36 (1733)** | 0.19 (720) | **0.36 (1353)** | 0.34 (937) | 0.26 (120) | 0.29 (258) |
| SDV | **0.45 (1733)** | 0.41 (1443) | **0.45 (1460)** | 0.26 (677) | **0.45 (536)** | 0.43 (1086) |
| 4W | 0.59 (1733) | 0.46 (1443) | 0.61 (1145) | 0.47 (1733) | **0.62 (494)** | 0.55 (1163) |
| 8W GH FT | **0.47 (1733)** | 0.46 (1220) | 0.46 (1188) | 0.43 (1598) | 0.31 (929) | 0.26 (952) |
| FT Field | 0.50 (1733) | 0.31 (911) | 0.46 (1300) | **0.52 (1514)** | 0.36 (1114) | 0.39 (1659) |
| FRI | 0.11 (1733) | 0.20 (335) | 0.21 (178) | **0.24 (58)** | 0.06 (557) | 0.11 (1005) |
| 0W GH FT | **0.64 (1733)** | 0.61 (1530) | 0.60 (1503) | 0.58 (1733) | 0.61 (932) | 0.60 (831) |
| FT GH | 0.55 (1733) | 0.54 (1194) | 0.55 (1313) | 0.54 (1733) | **0.59 (1194)** | 0.46 (1194) |
| SD | **0.62 (1733)** | 0.60 (1429) | 0.58 (790) | 0.56 (1733) | 0.57 (868) | 0.55 (1414) |
| 0W | 0.49 (1733) | 0.47 (1402) | 0.46 (1163) | 0.48 (1617) | 0.50 (635) | **0.54 (1456)** |
| LDV | 0.62 (1733) | 0.60 (1561) | 0.59 (1319) | **0.64 (1733)** | 0.54 (696) | 0.48 (1336) |
| FLC | **0.28 (1733)** | 0.18 (1243) | 0.20 (1339) | 0.22 (66) | 0.19 (468) | 0.13 (624) |
| 2W | 0.65 (1733) | 0.51 (1243) | 0.59 (1153) | 0.54 (1733) | **0.66 (787)** | 0.49 (851) |

Color Legend: 0.66, 0.06

Figure 7: The regression performances of SPADIS, SConES(S), SConES(R), Univariate, Lasso(R) and Lasso(S) in PSS2 for GM network and all phenotypes. Rows are phenotypes and the columns are compared methods. The numbers in each cell show Pearson's squared correlation coefficients for the corresponding phenotype and method. The cardinality of the selected SNP sets are given in parentheses. For each phenotype, the best performing method is shown with bold and red font.

**Pearson's squared correlation coefficient, GI**

| | SPADIS | SConES(S) | SConES(R) | Univariate | Lasso(R) | Lasso(S) |
|---|---|---|---|---|---|---|
| LN22 | **0.45 (1733)** | 0.35 (1404) | 0.39 (1241) | 0.34 (1733) | 0.38 (414) | 0.44 (826) |
| LN16 | **0.59 (1733)** | 0.57 (1469) | 0.53 (1393) | 0.53 (1733) | 0.57 (1248) | 0.41 (814) |
| LN10 | 0.45 (1733) | 0.41 (1356) | 0.48 (1291) | 0.42 (1733) | **0.49 (347)** | 0.35 (510) |
| 0W GH LN | 0.38 (1733) | 0.33 (1166) | 0.37 (1089) | 0.32 (360) | **0.39 (755)** | 0.35 (1274) |
| 8W GH LN | **0.37 (1733)** | 0.25 (1021) | 0.36 (1299) | 0.34 (937) | 0.26 (120) | 0.29 (258) |
| SDV | **0.46 (1733)** | 0.40 (1327) | 0.44 (1458) | 0.26 (677) | 0.45 (536) | 0.43 (1086) |
| 4W | 0.59 (1733) | 0.46 (1452) | 0.61 (1220) | 0.47 (1733) | **0.62 (494)** | 0.55 (1163) |
| 8W GH FT | **0.47 (1733)** | **0.47 (1288)** | 0.46 (1125) | 0.43 (1598) | 0.31 (929) | 0.26 (952) |
| FT Field | 0.50 (1733) | 0.32 (932) | 0.47 (1388) | **0.52 (1514)** | 0.36 (1114) | 0.39 (1659) |
| FRI | 0.11 (1733) | 0.21 (334) | 0.18 (199) | **0.24 (58)** | 0.06 (557) | 0.11 (1005) |
| 0W GH FT | **0.63 (1733)** | 0.61 (1582) | 0.60 (1481) | 0.58 (1733) | 0.61 (932) | 0.60 (831) |
| FT GH | 0.56 (1733) | 0.56 (1411) | 0.57 (1291) | 0.54 (1733) | **0.59 (1194)** | 0.46 (1194) |
| SD | **0.64 (1733)** | 0.60 (1435) | 0.56 (799) | 0.56 (1733) | 0.57 (868) | 0.55 (1414) |
| 0W | 0.50 (1733) | 0.49 (1514) | 0.44 (1124) | 0.48 (1617) | 0.50 (635) | **0.54 (1456)** |
| LDV | 0.63 (1733) | 0.59 (1576) | 0.59 (1218) | **0.64 (1733)** | 0.54 (696) | 0.48 (1336) |
| FLC | **0.28 (1733)** | 0.21 (1346) | 0.20 (1253) | 0.22 (66) | 0.19 (468) | 0.13 (624) |
| 2W | 0.64 (1733) | 0.62 (1425) | 0.61 (1191) | 0.54 (1733) | **0.66 (787)** | 0.49 (851) |

Color Legend: 0.66 – 0.06

Figure 8: The regression performances of SPADIS, SConES(S), SConES(R), Univariate, Lasso(R) and Lasso(S) in PSS2 for GI network and all phenotypes. Rows are phenotypes and the columns are compared methods. The numbers in each cell show Pearson's squared correlation coefficients for the corresponding phenotype and method. The cardinality of the selected SNP sets are given in parentheses. For each phenotype, the best performing method is shown with bold and red font.

**Pearson's squared correlation coefficient, GS-HICN**

| | SPADIS | SConES(S) | SConES(R) | Univariate | Lasso(R) | Lasso(S) |
|---|---|---|---|---|---|---|
| LN22 | **0.50 (1733)** | 0.35 (1323) | 0.40 (1252) | 0.34 (1733) | 0.38 (414) | 0.44 (826) |
| LN16 | **0.58 (1733)** | 0.55 (1386) | 0.53 (1272) | 0.53 (1733) | 0.57 (1248) | 0.41 (814) |
| LN10 | 0.48 (1733) | 0.44 (1453) | 0.45 (1180) | 0.42 (1733) | **0.49 (347)** | 0.35 (510) |
| 0W GH LN | **0.42 (1733)** | 0.30 (1159) | 0.34 (1000) | 0.32 (360) | 0.39 (755) | 0.35 (1274) |
| 8W GH LN | 0.35 (1733) | 0.27 (382) | **0.36 (1357)** | 0.34 (937) | 0.26 (120) | 0.29 (258) |
| SDV | 0.39 (1733) | 0.42 (1393) | 0.43 (1452) | 0.26 (677) | **0.45 (536)** | 0.43 (1086) |
| 4W | 0.58 (1733) | 0.47 (1360) | **0.63 (1095)** | 0.47 (1733) | 0.62 (494) | 0.55 (1163) |
| 8W GH FT | 0.41 (1733) | 0.47 (1183) | **0.48 (1204)** | 0.43 (1598) | 0.31 (929) | 0.26 (952) |
| FT Field | **0.52 (1733)** | 0.31 (1013) | 0.48 (1319) | **0.52 (1514)** | 0.36 (1114) | 0.39 (1659) |
| FRI | 0.13 (1733) | 0.20 (268) | 0.19 (239) | **0.24 (58)** | 0.06 (557) | 0.11 (1005) |
| 0W GH FT | **0.64 (1733)** | 0.61 (1557) | 0.62 (1388) | 0.58 (1733) | 0.61 (932) | 0.60 (831) |
| FT GH | 0.57 (1733) | 0.55 (1462) | 0.55 (1308) | 0.54 (1733) | **0.59 (1194)** | 0.46 (1194) |
| SD | **0.64 (1733)** | 0.60 (1464) | 0.57 (762) | 0.56 (1733) | 0.57 (868) | 0.55 (1414) |
| 0W | **0.54 (1733)** | 0.50 (1304) | 0.44 (923) | 0.48 (1617) | 0.50 (635) | **0.54 (1456)** |
| LDV | **0.65 (1733)** | 0.55 (974) | 0.60 (1224) | 0.64 (1733) | 0.54 (696) | 0.48 (1336) |
| FLC | **0.26 (1733)** | 0.17 (1322) | 0.17 (1269) | 0.22 (66) | 0.19 (468) | 0.13 (624) |
| 2W | 0.62 (1733) | 0.63 (1520) | 0.61 (1195) | 0.54 (1733) | **0.66 (787)** | 0.49 (851) |

Color Legend: 0.66 – 0.06

Figure 9: The regression performances of SPADIS, SConES(S), SConES(R), Univariate, Lasso(R) and Lasso(S) in PSS2 for GS-HICN network and all phenotypes. Rows are phenotypes and the columns are compared methods. The numbers in each cell show Pearson's squared correlation coefficients for the corresponding phenotype and method. The cardinality of the selected SNP sets are given in parentheses. For each phenotype, the best performing method is shown with bold and red font.

**Pearson's squared correlation coefficient**
**SPADIS - SConES(S), PSS2**

| | GS | GM | GI | GS-HICN |
|---|---|---|---|---|
| LN22 | **0.47 - 0.35*** | **0.45 - 0.35*** | **0.45 - 0.35*** | **0.50 - 0.35*** |
| LN16 | 0.57 - 0.56 | 0.58 - 0.57 | 0.59 - 0.57 | 0.58 - 0.55 |
| LN10 | 0.48 - 0.42 | 0.45 - 0.43 | 0.45 - 0.41 | 0.48 - 0.44 |
| 0W GH LN | **0.43 - 0.28*** | 0.39 - 0.29 | 0.38 - 0.33 | 0.42 - 0.30 |
| 8W GH LN | 0.36 - 0.24 | **0.36 - 0.19*** | 0.37 - 0.25 | 0.35 - 0.27 |
| SDV | 0.43 - 0.40 | 0.45 - 0.41 | 0.46 - 0.40 | 0.39 - 0.42 |
| 4W | 0.58 - 0.46 | 0.59 - 0.46 | 0.59 - 0.46 | 0.58 - 0.47 |
| 8W GH FT | 0.47 - 0.46 | 0.47 - 0.46 | 0.47 - 0.47 | 0.41 - 0.47 |
| FT Field | **0.49 - 0.31*** | **0.50 - 0.31*** | **0.50 - 0.32*** | **0.52 - 0.31*** |
| FRI | 0.11 - 0.21 | 0.11 - 0.20 | 0.11 - 0.21 | 0.13 - 0.20 |
| 0W GH FT | **0.66 - 0.60*** | 0.64 - 0.61 | 0.63 - 0.61 | 0.64 - 0.61 |
| FT GH | 0.58 - 0.53 | 0.55 - 0.54 | 0.56 - 0.56 | 0.57 - 0.55 |
| SD | 0.60 - 0.60 | 0.62 - 0.60 | 0.64 - 0.60 | 0.64 - 0.60 |
| 0W | 0.52 - 0.44 | 0.49 - 0.47 | 0.50 - 0.49 | 0.54 - 0.50 |
| LDV | 0.62 - 0.52 | 0.62 - 0.60 | 0.63 - 0.59 | 0.65 - 0.55 |
| FLC | **0.28 - 0.17*** | **0.28 - 0.18*** | 0.28 - 0.21 | 0.26 - 0.17 |
| 2W | 0.64 - 0.60 | **0.65 - 0.51*** | 0.64 - 0.62 | 0.62 - 0.63 |

Color Legend: 0.21 – 0 – -0.21

**Pearson's squared correlation coefficient**
**SPADIS - SConES(R), PSS2**

| | GS | GM | GI | GS-HICN |
|---|---|---|---|---|
| LN22 | **0.47 - 0.38*** | 0.45 - 0.41 | 0.45 - 0.39 | **0.50 - 0.40*** |
| LN16 | 0.57 - 0.53 | 0.58 - 0.52 | 0.59 - 0.53 | 0.58 - 0.53 |
| LN10 | 0.48 - 0.48 | 0.45 - 0.48 | 0.45 - 0.48 | 0.48 - 0.45 |
| 0W GH LN | 0.43 - 0.35 | 0.39 - 0.36 | 0.38 - 0.37 | 0.42 - 0.34 |
| 8W GH LN | 0.36 - 0.36 | 0.36 - 0.36 | 0.37 - 0.36 | 0.35 - 0.36 |
| SDV | 0.43 - 0.44 | 0.45 - 0.45 | 0.46 - 0.44 | 0.39 - 0.43 |
| 4W | 0.58 - 0.61 | 0.59 - 0.61 | 0.59 - 0.61 | 0.58 - 0.63 |
| 8W GH FT | 0.47 - 0.47 | 0.47 - 0.46 | 0.47 - 0.46 | 0.41 - 0.48 |
| FT Field | 0.49 - 0.49 | 0.50 - 0.46 | 0.50 - 0.47 | 0.52 - 0.48 |
| FRI | 0.11 - 0.22 | 0.11 - 0.21 | 0.11 - 0.18 | 0.13 - 0.19 |
| 0W GH FT | 0.66 - 0.61 | 0.64 - 0.60 | 0.63 - 0.60 | 0.64 - 0.62 |
| FT GH | 0.58 - 0.57 | 0.55 - 0.55 | 0.56 - 0.57 | 0.57 - 0.55 |
| SD | 0.60 - 0.58 | 0.62 - 0.58 | 0.64 - 0.56 | 0.64 - 0.57 |
| 0W | 0.52 - 0.45 | 0.49 - 0.46 | 0.50 - 0.44 | 0.54 - 0.44 |
| LDV | 0.62 - 0.58 | 0.62 - 0.59 | 0.63 - 0.59 | 0.65 - 0.60 |
| FLC | 0.28 - 0.20 | 0.28 - 0.20 | 0.28 - 0.20 | 0.26 - 0.17 |
| 2W | 0.64 - 0.61 | 0.65 - 0.59 | 0.64 - 0.61 | 0.62 - 0.61 |

Color Legend: 0.11 – 0 – -0.11

Figure 10: The regression performances of SPADIS, SConES(R) and SConES(S) on AT data for PSS2. Rows represent different phenotypes and columns represent different networks. The numbers in each cell show Pearson's squared correlation coefficients achieved by SPADIS and SConES, respectively (separated with a dash). The tone of background color reflects the difference in correlation coefficients between SPADIS and SConES. Red indicates SPADIS performs better than SConES while blue indicates otherwise. Differences that are found to be statistically significant are shown in bold and white and marked with star (*).

**Pearson's squared correlation coefficient**
**SPADIS - Lasso(S), PSS2**

| | GS | GM | GI | GS-HICN |
|---|---|---|---|---|
| LN22 | 0.47 - 0.44 | 0.45 - 0.44 | 0.45 - 0.44 | 0.50 - 0.44 |
| LN16 | 0.57 - 0.41* | 0.58 - 0.41* | 0.59 - 0.41* | 0.58 - 0.41* |
| LN10 | 0.48 - 0.35 | 0.45 - 0.35 | 0.45 - 0.35 | 0.48 - 0.35 |
| 0W GH LN | 0.43 - 0.35* | 0.39 - 0.35 | 0.38 - 0.35 | 0.42 - 0.35 |
| 8W GH LN | 0.36 - 0.29 | 0.36 - 0.29 | 0.37 - 0.29 | 0.35 - 0.29 |
| SDV | 0.43 - 0.43 | 0.45 - 0.43 | 0.46 - 0.43 | 0.39 - 0.43 |
| 4W | 0.58 - 0.55 | 0.59 - 0.55 | 0.59 - 0.55 | 0.58 - 0.55 |
| 8W GH FT | 0.47 - 0.26* | 0.47 - 0.26* | 0.47 - 0.26* | 0.41 - 0.26* |
| FT Field | 0.49 - 0.39 | 0.50 - 0.39* | 0.50 - 0.39* | 0.52 - 0.39* |
| FRI | 0.11 - 0.11 | 0.11 - 0.11 | 0.11 - 0.11 | 0.13 - 0.11 |
| 0W GH FT | 0.66 - 0.60* | 0.64 - 0.60 | 0.63 - 0.60 | 0.64 - 0.60 |
| FT GH | 0.58 - 0.46* | 0.55 - 0.46 | 0.56 - 0.46 | 0.57 - 0.46* |
| SD | 0.60 - 0.55 | 0.62 - 0.55 | 0.64 - 0.55* | 0.64 - 0.55* |
| 0W | 0.52 - 0.54 | 0.49 - 0.54 | 0.50 - 0.54 | 0.54 - 0.54 |
| LDV | 0.62 - 0.48* | 0.62 - 0.48* | 0.63 - 0.48* | 0.65 - 0.48* |
| FLC | 0.28 - 0.13* | 0.28 - 0.13* | 0.28 - 0.13* | 0.26 - 0.13* |
| 2W | 0.64 - 0.49* | 0.65 - 0.49* | 0.64 - 0.49* | 0.62 - 0.49 |

Color Legend: 0.22, 0, -0.22

**Pearson's squared correlation coefficient**
**SPADIS - Lasso(R), PSS2**

| | GS | GM | GI | GS-HICN |
|---|---|---|---|---|
| LN22 | 0.47 - 0.38 | 0.45 - 0.38 | 0.45 - 0.38 | 0.50 - 0.38* |
| LN16 | 0.57 - 0.57 | 0.58 - 0.57 | 0.59 - 0.57 | 0.58 - 0.57 |
| LN10 | 0.48 - 0.49 | 0.45 - 0.49 | 0.45 - 0.49 | 0.48 - 0.49 |
| 0W GH LN | 0.43 - 0.39 | 0.39 - 0.39 | 0.38 - 0.39 | 0.42 - 0.39 |
| 8W GH LN | 0.36 - 0.26 | 0.36 - 0.26 | 0.37 - 0.26 | 0.35 - 0.26 |
| SDV | 0.43 - 0.45 | 0.45 - 0.45 | 0.46 - 0.45 | 0.39 - 0.45 |
| 4W | 0.58 - 0.62 | 0.59 - 0.62 | 0.59 - 0.62 | 0.58 - 0.62 |
| 8W GH FT | 0.47 - 0.31* | 0.47 - 0.31* | 0.47 - 0.31* | 0.41 - 0.31 |
| FT Field | 0.49 - 0.36 | 0.50 - 0.36 | 0.50 - 0.36 | 0.52 - 0.36* |
| FRI | 0.11 - 0.06 | 0.11 - 0.06 | 0.11 - 0.06 | 0.13 - 0.06 |
| 0W GH FT | 0.66 - 0.61 | 0.64 - 0.61 | 0.63 - 0.61 | 0.64 - 0.61 |
| FT GH | 0.58 - 0.59 | 0.55 - 0.59 | 0.56 - 0.59 | 0.57 - 0.59 |
| SD | 0.60 - 0.57 | 0.62 - 0.57 | 0.64 - 0.57 | 0.64 - 0.57 |
| 0W | 0.52 - 0.50 | 0.49 - 0.50 | 0.50 - 0.50 | 0.54 - 0.50 |
| LDV | 0.62 - 0.54 | 0.62 - 0.54 | 0.63 - 0.54 | 0.65 - 0.54* |
| FLC | 0.28 - 0.19 | 0.28 - 0.19 | 0.28 - 0.19 | 0.26 - 0.19 |
| 2W | 0.64 - 0.66 | 0.65 - 0.66 | 0.64 - 0.66 | 0.62 - 0.66 |

Color Legend: 0.16, 0, -0.16

Figure 11: The regression performances of SPADIS, Lasso(R) and Lasso(S) on AT data for PSS2. Rows represent different phenotypes and columns represent different networks. The numbers in each cell show Pearson's squared correlation coefficients achieved by SPADIS and Lasso, respectively (separated with a dash). The tone of background color reflects the difference in correlation coefficients between SPADIS and Lasso. Red indicates SPADIS performs better than Lasso while blue indicates otherwise. Differences that are found to be statistically significant are shown in bold and white and marked with star (*).

**Pearson's squared correlation coefficient**
**SPADIS - Univariate, PSS2**

| | GS | GM | GI | GS-HICN |
|---|---|---|---|---|
| LN22 | 0.47 - 0.34* | 0.45 - 0.34* | 0.45 - 0.34* | 0.50 - 0.34* |
| LN16 | 0.57 - 0.53 | 0.58 - 0.53 | 0.59 - 0.53 | 0.58 - 0.53 |
| LN10 | 0.48 - 0.42 | 0.45 - 0.42 | 0.45 - 0.42 | 0.48 - 0.42 |
| 0W GH LN | 0.43 - 0.32 | 0.39 - 0.32 | 0.38 - 0.32 | 0.42 - 0.32 |
| 8W GH LN | 0.36 - 0.34 | 0.36 - 0.34 | 0.37 - 0.34 | 0.35 - 0.34 |
| SDV | 0.43 - 0.26* | 0.45 - 0.26* | 0.46 - 0.26* | 0.39 - 0.26* |
| 4W | 0.58 - 0.47* | 0.59 - 0.47* | 0.59 - 0.47* | 0.58 - 0.47* |
| 8W GH FT | 0.47 - 0.43 | 0.47 - 0.43 | 0.47 - 0.43 | 0.41 - 0.43 |
| FT Field | 0.49 - 0.52 | 0.50 - 0.52 | 0.50 - 0.52 | 0.52 - 0.52 |
| FRI | 0.11 - 0.24 | 0.11 - 0.24 | 0.11 - 0.24 | 0.13 - 0.24 |
| 0W GH FT | 0.66 - 0.58* | 0.64 - 0.58 | 0.63 - 0.58 | 0.64 - 0.58 |
| FT GH | 0.58 - 0.54 | 0.55 - 0.54 | 0.56 - 0.54 | 0.57 - 0.54 |
| SD | 0.60 - 0.56 | 0.62 - 0.56 | 0.64 - 0.56 | 0.64 - 0.56* |
| 0W | 0.52 - 0.48 | 0.49 - 0.48 | 0.50 - 0.48 | 0.54 - 0.48 |
| LDV | 0.62 - 0.64 | 0.62 - 0.64 | 0.63 - 0.64 | 0.65 - 0.64 |
| FLC | 0.28 - 0.22 | 0.28 - 0.22 | 0.28 - 0.22 | 0.26 - 0.22 |
| 2W | 0.64 - 0.54* | 0.65 - 0.54* | 0.64 - 0.54* | 0.62 - 0.54 |

Color Legend: 0.20, 0, -0.20

Figure 12: The regression performances of SPADIS and Univariate on AT data for PSS2. Rows represent different phenotypes and columns represent different networks. The numbers in each cell show Pearson's squared correlation coefficients achieved by SPADIS and Univariate, respectively (separated with a dash). The tone of background color reflects the difference in correlation coefficients between SPADIS and Univariate. Red indicates SPADIS performs better than Univariate while blue indicates otherwise. Differences that are found to be statistically significant are shown in bold and white and marked with star (*).

**Pearson's squared correlation coefficient**
**GS-HICN - GS, k = 100**

| | SPADIS | SConES(R) | SConES(S) |
|---|---|---|---|
| LN22 | 0.33 - 0.33 | 0.37 - 0.37 | 0.30 - 0.31 |
| LN16 | 0.40 - 0.46 | 0.43 - 0.44 | 0.47 - 0.42 |
| LN10 | 0.29 - 0.28 | 0.22 - 0.20 | 0.29 - 0.21 |
| 0W GH LN | 0.20 - 0.28 | 0.28 - 0.28 | 0.18 - 0.20 |
| 8W GH LN | 0.22 - 0.29 | 0.26 - 0.28 | 0.16 - 0.19 |
| SDV | 0.21 - 0.33 | 0.16 - 0.17 | 0.21 - 0.16 |
| 4W | 0.53 - 0.42 | 0.40 - 0.39 | 0.44 - 0.43 |
| 8W GH FT | 0.41 - 0.38 | 0.35 - 0.37 | 0.32 - 0.34 |
| FT Field | 0.30 - 0.29 | 0.26 - 0.23 | 0.27 - 0.23 |
| FRI | 0.13 - 0.08 | 0.10 - 0.08 | 0.07 - 0.07 |
| 0W GH FT | 0.37 - 0.34 | 0.34 - 0.35 | 0.35 - 0.33 |
| FT GH | 0.44 - 0.48 | 0.42 - 0.39 | 0.43 - 0.37 |
| SD | 0.52 - 0.58 | 0.56 - 0.58 | 0.57 - 0.55 |
| 0W | 0.30 - 0.34 | 0.23 - 0.21 | 0.27 - 0.25 |
| LDV | 0.42 - 0.42 | 0.44 - 0.44 | 0.40 - 0.45 |
| FLC | 0.22 - 0.13 | 0.10 - 0.06 | 0.08 - 0.07 |
| 2W | 0.40 - 0.39 | 0.28 - 0.30 | 0.34 - 0.31 |

Color Legend: 0.12, 0, -0.12

**Pearson's squared correlation coefficient**
**GS-HICN - GM, k = 100**

| | SPADIS | SConES(R) | SConES(S) |
|---|---|---|---|
| LN22 | 0.33 - 0.36 | 0.37 - 0.37 | 0.30 - 0.29 |
| LN16 | 0.40 - 0.44 | 0.43 - 0.44 | 0.47 - 0.43 |
| LN10 | 0.29 - 0.28 | 0.22 - 0.21 | 0.29 - 0.23 |
| 0W GH LN | 0.20 - 0.29 | 0.28 - 0.29 | 0.18 - 0.23 |
| 8W GH LN | 0.22 - 0.31 | 0.26 - 0.27 | 0.16 - 0.21 |
| SDV | 0.21 - 0.34 | 0.16 - 0.17 | 0.21 - 0.15 |
| 4W | 0.53 - 0.34 | 0.40 - 0.41 | 0.44 - 0.44 |
| 8W GH FT | 0.41 - 0.42 | 0.35 - 0.37 | 0.32 - 0.32 |
| FT Field | 0.30 - 0.31 | 0.26 - 0.23 | 0.27 - 0.25 |
| FRI | 0.13 - 0.07 | 0.10 - 0.10 | 0.07 - 0.06 |
| 0W GH FT | 0.37 - 0.35 | 0.34 - 0.35 | 0.35 - 0.32 |
| FT GH | 0.44 - 0.41 | 0.42 - 0.39 | 0.43 - 0.39 |
| SD | 0.52 - 0.55 | 0.56 - 0.58 | 0.57 - 0.57 |
| 0W | 0.30 - 0.27 | 0.23 - 0.20 | 0.27 - 0.23 |
| LDV | 0.42 - 0.46 | 0.44 - 0.46 | 0.40 - 0.52 |
| FLC | 0.22 - 0.12 | 0.10 - 0.06 | 0.08 - 0.06 |
| 2W | 0.40 - 0.32 | 0.28 - 0.31 | 0.34 - 0.33 |

Color Legend: 0.19, 0, -0.19

**Pearson's squared correlation coefficient**
**GS-HICN - GI, k = 100**

| | SPADIS | SConES(R) | SConES(S) |
|---|---|---|---|
| LN22 | 0.33 - 0.32 | 0.37 - 0.36 | 0.30 - 0.31 |
| LN16 | 0.40 - 0.44 | 0.43 - 0.44 | 0.47 - 0.45 |
| LN10 | 0.29 - 0.30 | 0.22 - 0.20 | 0.29 - 0.26 |
| 0W GH LN | 0.20 - 0.33 | 0.28 - 0.24 | 0.18 - 0.17 |
| 8W GH LN | 0.22 - 0.29 | 0.26 - 0.28 | 0.16 - 0.22 |
| SDV | 0.21 - 0.34 | 0.16 - 0.20 | 0.21 - 0.18 |
| 4W | 0.53 - 0.37 | 0.40 - 0.41 | 0.44 - 0.36 |
| 8W GH FT | 0.41 - 0.41 | 0.35 - 0.36 | 0.32 - 0.35 |
| FT Field | 0.30 - 0.32 | 0.26 - 0.25 | 0.27 - 0.21 |
| FRI | 0.13 - 0.11 | 0.10 - 0.10 | 0.07 - 0.11 |
| 0W GH FT | 0.37 - 0.38 | 0.34 - 0.35 | 0.35 - 0.33 |
| FT GH | 0.44 - 0.44 | 0.42 - 0.35 | 0.43 - 0.41 |
| SD | 0.52 - 0.53 | 0.56 - 0.55 | 0.57 - 0.56 |
| 0W | 0.30 - 0.20 | 0.23 - 0.22 | 0.27 - 0.27 |
| LDV | 0.42 - 0.46 | 0.44 - 0.44 | 0.40 - 0.43 |
| FLC | 0.22 - 0.12 | 0.10 - 0.06 | 0.08 - 0.06 |
| 2W | 0.40 - 0.37 | 0.28 - 0.29 | 0.34 - 0.27 |

Color Legend: 0.16, 0, -0.16

Figure 13: This figure shows the effect of using Hi-C data on the regression performances of SPADIS, SConES(R) and SConES(S) as compared to other networks. AT data is used with PSS1 and $k = 100$. Rows are the phenotypes and columns are the methods. The Pearson's squared correlation coefficient differences between GS-HICN and other networks are shown: GS-HICN vs GS (left), GS-HICN vs GM (center), and GS-HICN vs GI (right). The tone of background color reflects the difference in obtained Pearson's correlation coefficients between using GS-HICN or other networks. Red indicates GS-HICN performs better while blue indicates otherwise. Differences that are found to be statistically significant are shown in bold and white and marked with star (*).

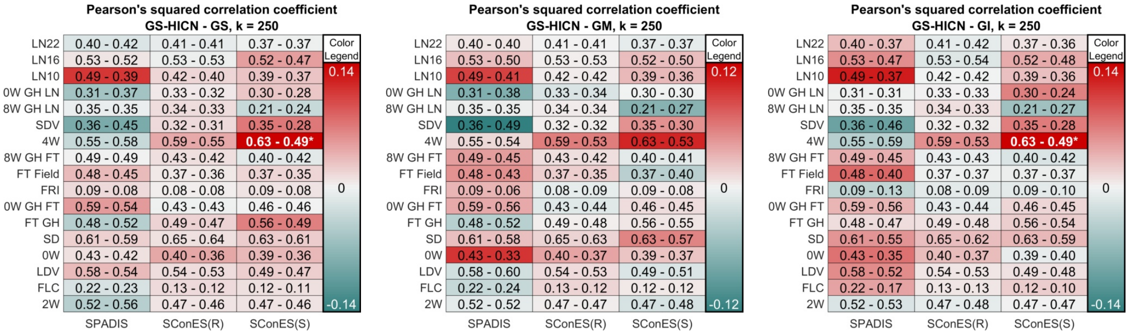


Figure 14: This figure shows the effect of using Hi-C data on the regression performances of SPADIS, SConES(R) and SConES(S) as compared to other networks. AT data is used with PSS1 and $k = 250$. Rows are the phenotypes and columns are the methods. The Pearson's squared correlation coefficient differences between GS-HICN and other networks are shown: GS-HICN vs GS (left), GS-HICN vs GM (center), and GS-HICN vs GI (right). The tone of background color reflects the difference in obtained Pearson's correlation coefficients between using GS-HICN or other networks. Red indicates GS-HICN performs better while blue indicates otherwise. Differences that are found to be statistically significant are shown in bold and white and marked with star (*).

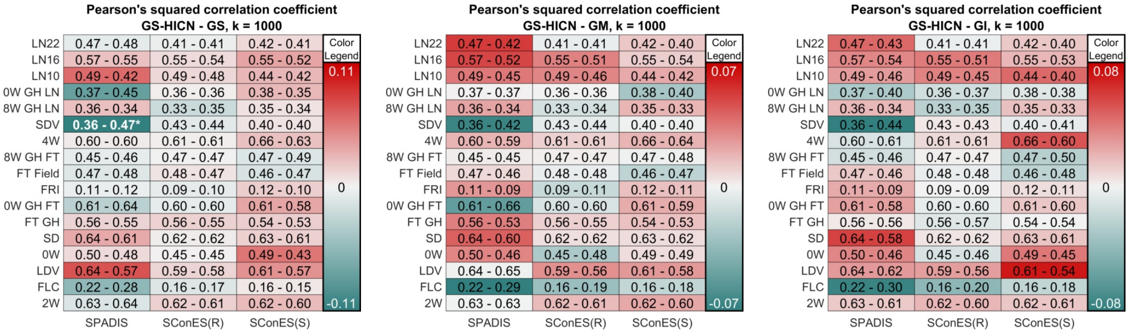


Figure 15: This figure shows the effect of using Hi-C data on the regression performances of SPADIS, SConES(R) and SConES(S) as compared to other networks. AT data is used with PSS1 and $k = 1000$. Rows are the phenotypes and columns are the methods. The Pearson's squared correlation coefficient differences between GS-HICN and other networks are shown: GS-HICN vs GS (left), GS-HICN vs GM (center), and GS-HICN vs GI (right). The tone of background color reflects the difference in obtained Pearson's correlation coefficients between using GS-HICN or other networks. Red indicates GS-HICN performs better while blue indicates otherwise. Differences that are found to be statistically significant are shown in bold and white and marked with star (*).

### 2 Supplementary Tables

Table 1: Table shows statistics about the genes and functionalities that are hit by the selected SNP sets by each method for PSS1 and $k = 100$. All statistics given are averaged over all 17 phenotypes. GenesHit is the number of distinct candidate flowering time genes identified. GO-Hit is the number of distinct biological processes hit by hitting an associated gene with that GO term. Precision is the ratio of number of selected SNPs near candidate genes and total number of selected SNPs. $R^2$ is the Pearson's squared correlation coefficient.

| Network | Method | Genes-Hit | GO-Hit | Precision(%) | $R^2$ |
|---|---|---|---|---|---|
| GS | SPADIS | 6.2 | 156 | 7.5% | 0.343 |
| GS | SConES(S) | 4.2 | 107 | 11.4% | 0.287 |
| GS | SConES(R) | 4.4 | 114 | 11.2% | 0.302 |
| GM | SPADIS | 6.3 | 153 | 7.3% | 0.331 |
| GM | SConES(S) | 4.5 | 116 | 11.2% | 0.297 |
| GM | SConES(R) | 4.6 | 118 | 11.0% | 0.306 |
| GI | SPADIS | 4.8 | 139 | 6.2% | 0.336 |
| GI | SConES(S) | 4.0 | 114 | 10.4% | 0.291 |
| GI | SConES(R) | 4.3 | 116 | 10.4% | 0.301 |
| GS-HICN | SPADIS | 6.1 | 157 | 7.2% | 0.335 |
| GS-HICN | SConES(S) | 4.7 | 118 | 11.1% | 0.302 |
| GS-HICN | SConES(R) | 4.7 | 119 | 10.9% | 0.306 |

Table 2: Table shows statistics about the genes and functionalities that are hit by the selected SNP sets by each method for PSS1 and $k = 250$. All statistics given are averaged over all 17 phenotypes. GenesHit is the number of distinct candidate flowering time genes identified. GO-Hit is the number of distinct biological processes hit by hitting an associated gene with that GO term. Precision is the ratio of number of selected SNPs near candidate genes and total number of selected SNPs. $R^2$ is the Pearson's squared correlation coefficient.

| Network | Method | Genes-Hit | GO-Hit | Precision(%) | $R^2$ |
|---|---|---|---|---|---|
| GS | SPADIS | 13.8 | 312 | 6.8% | 0.441 |
| GS | SConES(S) | 8.3 | 213 | 9.9% | 0.371 |
| GS | SConES(R) | 8.9 | 232 | 9.6% | 0.396 |
| GM | SPADIS | 13.7 | 308 | 6.5% | 0.432 |
| GM | SConES(S) | 9.1 | 237 | 9.6% | 0.388 |
| GM | SConES(R) | 9.1 | 237 | 9.4% | 0.399 |
| GI | SPADIS | 10.0 | 289 | 5.1% | 0.415 |
| GI | SConES(S) | 7.8 | 228 | 8.4% | 0.376 |
| GI | SConES(R) | 8.5 | 235 | 8.9% | 0.402 |
| GS-HICN | SPADIS | 14.1 | 317 | 6.8% | 0.440 |
| GS-HICN | SConES(S) | 9.6 | 240 | 9.8% | 0.397 |
| GS-HICN | SConES(R) | 9.3 | 240 | 9.5% | 0.408 |

Table 3: Table shows statistics about the genes and functionalities that are hit by the selected SNP sets by each method for PSS1 and $k = 1000$. All statistics given are averaged over all 17 phenotypes. GenesHit is the number of distinct candidate flowering time genes identified. GO-Hit is the number of distinct biological processes hit by hitting an associated gene with that GO term. Precision is the ratio of number of selected SNPs near candidate genes and total number of selected SNPs. $R^2$ is the Pearson's squared correlation coefficient.

| Network | Method | Genes-Hit | GO-Hit | Precision(%) | $R^2$ |
|---|---|---|---|---|---|
| GS | SPADIS | 42.1 | 771 | 6.1% | 0.479 |
| GS | SConES(S) | 21.8 | 540 | 8.0% | 0.447 |
| GS | SConES(R) | 24.2 | 584 | 7.8% | 0.459 |
| GM | SPADIS | 42.5 | 733 | 6.4% | 0.466 |
| GM | SConES(S) | 25.0 | 596 | 7.9% | 0.459 |
| GM | SConES(R) | 25.0 | 601 | 7.7% | 0.459 |
| GI | SPADIS | 35.0 | 697 | 6.3% | 0.467 |
| GI | SConES(S) | 20.8 | 580 | 6.1% | 0.452 |
| GI | SConES(R) | 23.7 | 596 | 7.1% | 0.458 |
| GS-HICN | SPADIS | 43.5 | 789 | 6.5% | 0.474 |
| GS-HICN | SConES(S) | 26.8 | 609 | 7.9% | 0.466 |
| GS-HICN | SConES(R) | 25.9 | 607 | 7.8% | 0.460 |

### 3 Supplementary Text

#### 3.1 Parameter Selection

In both PSS1 and PSS2, measurements of regression performance are done by applying ridge regression in a 10-fold cross-validated setting. In PSS2, due to memory constraints, a parameter value that results in SNP sets with cardinality $> 3466$ ($1733 \times 2$) in at least one training fold are deemed invalid.
In PSS1, parameter selection for each method is performed as follows:

**SPADIS**: The distance parameter ($D$) and the penalty parameter ($\beta$) are selected by applying two consecutive line searches. First, seven different values of $D$ within $[D_{min}, D_{max}]$ varying in logarithmic scale are tested when $\beta = \infty$ and the $D$ value that maximizes the training regression performance is selected. $D_{min}$ is the minimum edge weight i.e. $D_{min} = \min_{e \in E} w(e)$. $D_{max}$ is the maximum distance such that penalty can be avoided i.e. $K(i, j) = 0$, $\forall i, j \in S^*$ where $S^*$ is the selected set when $\beta = \infty$ and $D = D_{max}$. We find $D_{max}$ by binary search. Having set the $D$ value, 16 $\beta$ values within $[10^{-4}, 1]\beta_{max}$ in logarithmic scale are tested where $\beta_{max} = 2kD \max(\mathbf{c})$. The $\beta$ value that maximizes the training regression performance is selected.

**SConES(S)**: We perform a line search for the connectivity parameter ($\lambda$) within the range $\left[\frac{\min(\mathbf{c})}{\delta}, \frac{\max(\mathbf{c})}{\delta}\right]$, in logarithmic scale where $\delta$ is the average degree of the graph. For each $\lambda$ value, the sparsity parameter ($\eta$) is set such that the number of selected SNPs is close to $k$. This is achieved by a binary search. Then, the most *stable* $\lambda$ is selected using the stability criteria [2]. Note that this is the parameter selection method described in [1]. We call the SConES runs with this parameter selection technique SConES(S) to distinguish it from the following version.

**SConES(R)**: Since, we select the parameters of SPADIS with respect to regression performance, for a fair comparison, we select SConES' parameters with respect to regression performance as well and call the runs of SConES with this technique SConES(R). As in SConES(S), we perform line search for $\lambda$ and binary search for $\eta$ parameters. The $\lambda$ value that maximizes training regression performance is selected.

**Lasso, GroupLasso and GraphLasso**: The regularization parameter ($\lambda$) is determined by binary search such that the number of selected SNPs is close to $k$.

In PSS2, parameter selection is performed as follows:
**SPADIS**: Same as done in PSS1. This time the number of SNPs to be selected ($k$) is set to 1733.

**SConES(S)**: First, binary search targeting 1733 is done on $\eta$ to find a valid lower bound for

$\eta$. Then, $7x7$ grid search experiments are conducted and the most *stable* parameter pair across the training folds is selected.

**SConES(R)**: First, a valid lower bound (that yields 1733 SNPs selected) for $\eta$ is found using binary search. Then, $7x7$ grid search experiments are conducted and the parameter pair that performs best in regression is selected.

**Lasso(S)**: First the lower bound of the regularization parameter ($\lambda$) that yields 1733 SNPs is found using binary search. Then, seven values of $\lambda$ in logarithmic scale are tested and the most *stable* one is selected.

**Lasso(R)**: First the lower bound of the regularization parameter ($\lambda$) that yields 1733 SNPs is found using binary search. Then, seven values of $\lambda$ in logarithmic scale are tested and the one that maximizes the regression performance on training data is selected. $\lambda$ values that result in SNP sets with cardinality $> 3466$ in at least one training fold are deemed invalid.
